# The trichothecene mycotoxin deoxynivalenol facilitates cell-to-cell invasion during wheat-tissue colonisation by *Fusarium graminearum*

**DOI:** 10.1101/2023.12.05.570169

**Authors:** Victoria Armer, Martin Urban, Tom Ashfield, Michael J. Deeks, Kim E. Hammond-Kosack

## Abstract

Fusarium Head Blight (FHB) disease on small grain cereals is primarily caused by the ascomycete fungal pathogen *Fusarium graminearum.* Infection of floral spike tissues is characterised by the biosynthesis and secretion of potent trichothecene mycotoxins, of which deoxynivalenol (DON) is widely reported due to its negative impacts on grain quality and consumer safety. The *TRI5* gene encodes an essential enzyme in the DON biosynthesis pathway and the single gene deletion mutant, *ΔTri5*, is widely reported to restrict disease progression to the inoculated spikelet. In this study, we present novel bioimaging evidence revealing that DON facilitates the traversal of the cell wall through plasmodesmata, a process essential for successful colonisation of host tissue. Chemical complementation of *ΔTri5* did not restore macro- or microscopic phenotypes, indicating that DON secretion is tightly regulated both spatially and temporally. A comparative qualitative and quantitative morphological cellular analysis revealed infections had no impact on plant cell wall thickness. Immuno- labelling of callose at plasmodesmata during infection indicates that DON can increase deposits when applied exogenously, but is reduced when *F. graminearum* hyphae are present. This study highlights the complexity of the inter-connected roles of mycotoxin production, cell wall architecture and plasmodesmata in this highly specialised interaction.

## Introduction

*Fusarium graminearum* (teleomorph *Gibberella zeae*) is an ascomycete fungal pathogen and the main causative agent of Fusarium Head Blight (FHB), or scab disease, on wheat. *F. graminearum* infects wheat floral tissues at flowering (anthesis), secreting many cell wall-degrading enzymes (CWDEs), other proteins and metabolites as well as mycotoxins that contaminate the developing grain, rendering it unsuitable for both human and livestock consumption (McMullen et al., 2012). Among these mycotoxins, the sesquiterpenoid type B toxins of the trichothecene class are particularly potent and include deoxynivalenol (DON), nivalenol (NIV), zearalenone (ZEA), and T-2 toxin (reviewed McCormick et al., 2011), all of which target the ribosome and inhibit protein synthesis (Brown et al., 2004). Trichothecene contamination of grain causes significant economic losses annually (McMullen et al., 1997), destroying wheat crops weeks before harvest and subsequently proliferating during ineffective grain storage/shipment. Epidemics of FHB occur when warm, wet weather coincides with anthesis and are particularly prominent in the mid-West USA, Asia, Brazil and Northern Europe (McMullen et al., 1997; Vaughan, Blackhouse and Del Ponte, 2016). Novel genetic targets are required to help control outbreaks of FHB disease due to the prevalence of resistance to the major class of azole fungicides in global *F. graminearum* strains (Fan et al., 2013). Incidences of FHB outbreaks are expected to increase as climate change increases precipitation around wheat harvests (Vaughan, Backhouse and Ponte, 2016). Hence, it is imperative that the infection biology of *F. graminearum* is explored further to aid in the development of resistant wheat varieties and precise chemical control, with the overall aim of minimising FHB-associated reductions in cereal yields, grain quality and to improve human/animal health.

The infection cycle of FHB commences with the dispersal of conidia (asexual) or ascospores (sexual) by rain droplet-induced splashes or wind onto wheat plants. During a typical infection of wheat at crop anthesis, germinating spores enter the host floral tissues through natural openings, such as stomata (Pritsch et al., 2000) and cracked open anther sacs, or have been reported to form penetration pegs on the abaxial surface of the palea and lemma tissues of the wheat spikelet (Wanjiru, Zhensheng and Buchenauer, 2002). Host-tissue colonisation continues with the invasive hyphae growing both intercellularly and intracellularly. *F. graminearum* has been noted to have a ‘biphasic’ lifestyle, whereby the advancing infection front is split between macroscopically symptomatic and symptomless phases (Brown et al., 2011). The symptomless phase is hallmarked by apoplastic growth, and the symptomatic by extensive intracellular growth. What initiates this switch is not yet known and is a subject of great interest. During later stages of infection, *F. graminearum* secretes CWDEs in abundance (Brown, Antoniw, and Hammond-Kosack, 2012) to facilitate infection by deconstructing wheat cell walls. At the rachis internode, invasive hyphae have been reported to enter vascular elements (Wanjiru et al., 2002) and grow through the remaining wheat spike within the vasculature as well as in the cortical tissue surrounding the vascular bundles (Brown et al., 2010). Furthermore, within the chlorenchyma band of the rachis, *F. graminearum* produces perithecia, sexual reproductive structures, completing its lifecycle (Guenther and Trail, 2005). Post-harvest, *F. graminearum* overwinters saprophytically on crop debris or within the soil, thereby infecting subsequent crop cycles. The presence of *F. graminearum* in the soil can be the primary cause of seedling blight and root rot in subsequent wheat crops (Parry et al., 1995).

Intracellular growth by *F. graminearum* has been previously reported to traverse wheat cell walls though pits, or pit fields, where plasmodesmata are present (Guenther and Trail, 2005; Jansen et al., 2005; Brown et al., 2010). PD are cytoplasmic communication channels that symplastically bridge the cell walls by an appressed endoplasmic reticulum (ER), known as a desmotubule, within a plasma membrane (PM) continuum stabilised by proteins connected to both the ER and PM (Sager and Lee, 2018). PD are instrumental to cellular signalling, allowing for the transport of sugars, ions and small proteins, to name a few. However, plants can adjust the permeability of PD by the deposition of callose, mediated by the action of callose synthases and β-1,3-glucanases (Lee and Lu, 2011) at PD junctions. This callose plugging leads to the symplastic isolation of cells that are damaged or under pathogen attack thereby restricting the movement of secreted pathogen effector proteins, toxins and other metabolites. PD have a major role in host plant defence against viruses, bacteria and fungi (Lee and Lu, 2011). *F. graminearum* exploits the plasmodesmatal transit highways by excreting β-1,3- glucanases: enzymes that catalyse the breakdown of the 1,3-O-glycosidic bond between glucose molecules in callose. RNA-seq analysis of *F. graminearum* infection of wheat spikes found that several Fusarium β-1,3-glucanases are upregulated in the host plant from as early as 6 hours post-infection and peaking between 36-48 hours after inoculation (Pritsch et al., 2007).

The trichothecene mycotoxin DON is a well-reported virulence factor in wheat floral tissues (Proctor et al., 1995; Jansen et al., 2005, Cuzick, Urban and Hammond-Kosack, 2008) and biosynthesis of the toxin requires the *TRI5* gene, encoding the enzyme trichothecene synthase (Hohn et al., 1993). DON is synthesised and then secreted by *F. graminearum* hyphae during infection and is a potent ribosomal-binding translational inhibitor. This broad-spectrum dampening of induced protein dependent defence responses has thus far prevented the elucidation of specific components of host immunity that restrict *F. graminearum* in the DON-deficient interaction. The low molecular weight of trichothecenes and their water-solubility allow them to rapidly enter cells and target eukaryotic ribosomes. This causes what is known as the ‘ribotoxic stress response’ that can activate, among other processes, mitogen-activated protein kinases (MAPKs) and apoptosis (reviewed Pestka, 2008). Deletion of *TRI5* eliminates the ability of *F. graminearum* to synthesise DON (Proctor et al., 1995), and infection of wheat floral tissues by the single gene deletion mutant (*ΔTri5*) is restricted to the inoculated spikelet, and results in the production of eye-shaped lesions on the outer glume (Jansen et al., 2005; Cuzick, Urban and Hammond-Kosack, 2008). Conversely, expression of *TRI5* in wild-type *F. graminearum* is correlated with DON accumulation *in planta* (Hallen-Adams et al., 2011). In non-host pathosystems, such as the model plant species *Arabidopsis thaliana*, infection of floral tissues with the single gene deletion mutant *ΔTri5* causes a wild-type disease phenotype, indicating that DON is not a virulence factor in this interaction (Cuzick, Urban and Hammond-Kosack, 2008). Current evidence indicates that the *TRI4* gene, which encodes a multi-functional cytochrome P450 monooxygenase and resides within the main trichothecene mycotoxin biosynthetic cluster (Tokai et al., 2007), is expressed during the *F. graminearum-*wheat coleoptile interaction (Qui et al., 2019), but the role of *TRI4* or *TRI5* as a virulence factors in coleoptiles has not yet been reported. The TRI4 protein is required for four consecutive oxygenation steps in trichothecene mycotoxin biosynthesis (Tokai et al., 2007), thus indicating a role for trichothecene mycotoxins during coleoptile infection. Through the use of fluorescent marker reporter strains, the *TRI5* gene has been shown to be induced during infection structure formation on wheat palea (Boenisch and Schäfer, 2011). However, the absence of *TRI5* in a *F. graminearum ΔTri5-GFP* strain did not impact the ability of *F. graminearum* to form infection cushions during initial time points of infection (Boenisch and Schäfer, 2011). The DON mycotoxin naturally occurs as two chemotypes, 15-ADON and 3-ADON, and individual *F. graminearum* strains secrete either toxin type. The wild-type (WT) strain used in this study, PH-1, synthesises 15-ADON. Host-plant resistance to DON is a characteristic of type II FHB resistance, whereby fungal advancement does not proceed beyond the rachis node (reviewed Mesterházy, 1995).

Whilst the macro-biology and some aspects of the cellular biology of the single-gene deletion mutant *ΔTri5* have been previously studied, the mode of restriction of *ΔTri5* remains to be elucidated. Postulations have been made around the role of DON during host-tissue colonisation, specifically relating to the targeting of ribosomes and the subsequent, broad-spectrum, protein translation inhibition (Pestka, 2010). However, what host defence mechanisms are targeted/specifically affected by DON have not been explored *in planta*. This study aims to re-evaluate the infection biology of the *ΔTri5* strain, and hence the role(s) of DON, during host-tissue colonisation through a combination of molecular and microscopy techniques. Through qualitative and quantitative image analysis of wheat floral tissues during WT and *ΔTri5* infection, we report that the *ΔTri5* single gene deletion mutant has an impaired ability to traverse plasmodesmata. We also find no evidence to support the hypothesis that a general increase in plant cell wall thickening occurs in the absence of DON production, whereby the upregulation of cell wall defences occurs during pathogen attack. From the data gathered, we infer that the secretion of DON during host-tissue colonisation is highly specific both spatially and temporally. This is indicated by the lack of increase in virulence in the *ΔTri5* mutant when supplied with DON at the point of inoculation in our study. In light of these discoveries, we pose new questions surrounding *F. graminearum* infection biology, cell wall colonisation and wheat host defence mechanisms.

## Results

The role(s) of DON during *F. graminearum* infection of different wheat tissues was addressed through a multifaceted approach. We applied a combination of detailed cell and molecular biology and morphological analyses of floral and coleoptile infections to analyse the effect of DON on hyphae traversing cell walls at PD and the occurrence of the defence response, callose deposition, at plasmodesmata during infections cause by either the wildtype (WT) strain or the single gene deletion mutant *ΔTri5 F. graminearum* strain.

### DON is not required for virulence on wheat coleoptiles and chemical complementation does not restore the WT disease phenotype on wheat spikes

To determine whether DON is or is not required for virulence on wheat coleoptiles under our conditions the fully susceptible cv. Apogee was tested. Inoculation of wheat coleoptiles revealed no differences in lesion length between the WT PH-1 strain and single gene deletion mutant *ΔTri5* (Fig. 1 (a) and 1(b)). However, RT-qPCR analysis showed that the WT strain expressed TRI5 during coleoptile infection at 3, S5 and 7 days post inoculation (dpi), but remains at very low levels (x0.8 FgActin) and stationary throughout the infection progression (Fig. 1(c)), indicating that TRI5 mRNA levels are not temporally altered for this interaction during the early to mid-time points explored. This finding supports a previous study by Qui et al. (2019), who reported accumulation of transcripts of the *TRI4* gene, also required for trichothecene mycotoxin biosynthesis.

**Figure 1.**
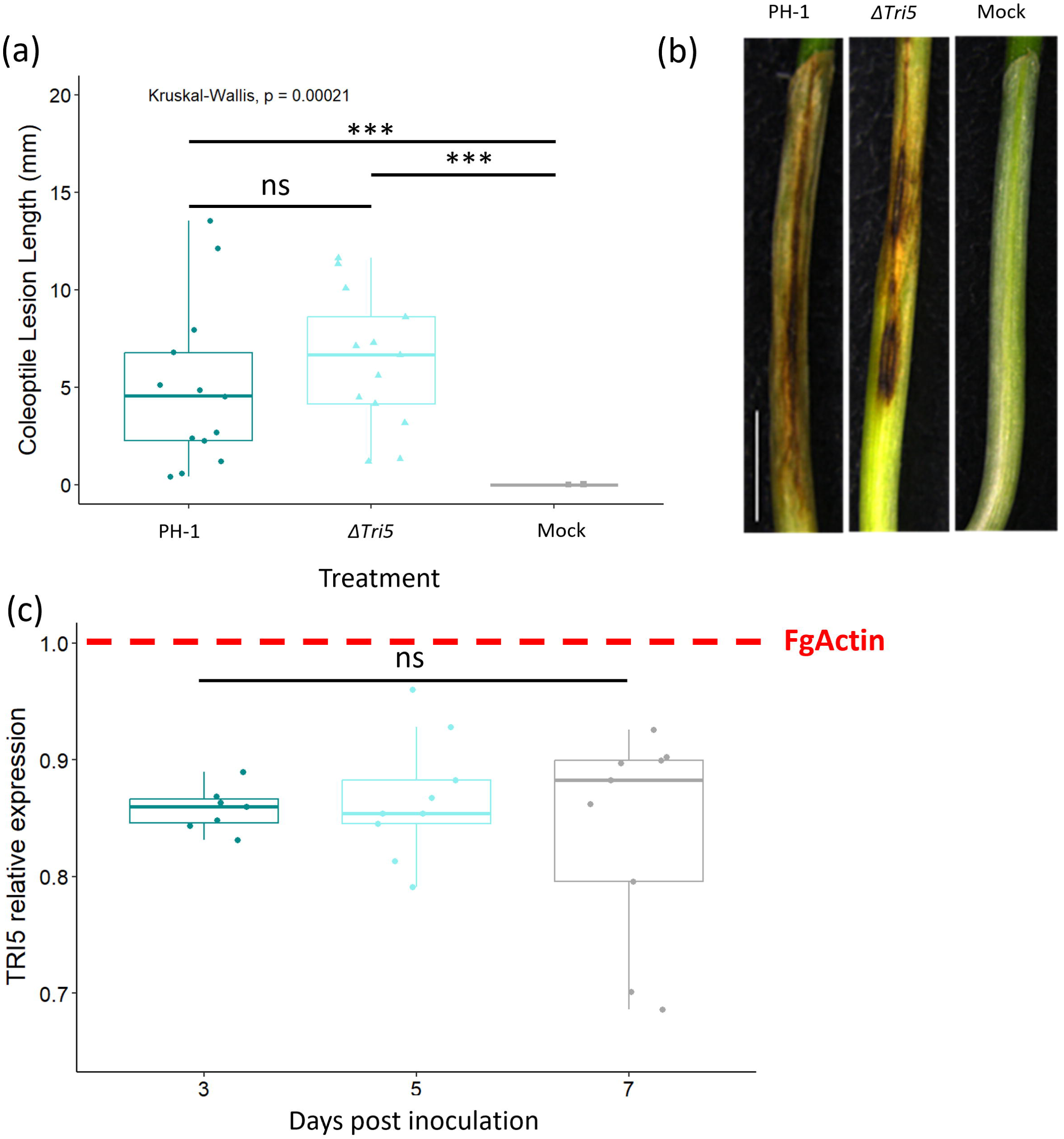
*F. graminearum* disease formation on wheat coleoptiles. (a) Length lesion at 7dpi for PH-1, the *ΔTri5* mutant and mock inoculations, Kruskal-Wallis *p* < 0.005(***). (b) Examples of disease lesion phenotypes at 7dpi for PH-1, *ΔTri5* and mock inoculations from rep 2, scale bar = 20mm and (c) Relative expression of *TRI5* measured using RT-qPCR at 3, 5 and 7dpi in wheat coleoptiles, normalised against FgActin expression. ANOVA F(2,22) = 0.421, p = 0.662 (ns).

Next, we asked whether the same host and pathogen genotypes showed different DON dependencies during floral tissue interactions. Disease progression of WT, *ΔTri5* and DON-complemented strains were analysed by tracking visible disease symptom development on the outer glume and rachis of inoculated wheat spikes. The single *ΔTri5* mutant was restricted to the inoculated spikelet in all instances. Whereas chemical complementation of the *ΔTri5* mutant with DON (35ppm) applied along with the conidia failed to restore the macroscopic WT spikelet phenotype occurring on the inoculated spikelet or spikelet-to-spikelet symptom development. This DON concentration was not detrimental to either spore germination or early spore germling growth (supplementary S1). Interestingly, co-inoculation of WT *F. graminearum* with DON at the same concentration did not result in any observable advancement of disease symptoms (Fig. 2 (a)). Application of DON (35ppm) alone did not induce any macroscopic disease symptoms and visually equated to the water only (dH_2_O) mock-inoculated samples (Fig. 2 (d), 3(a)). The area under the disease progression curve (AUDPC) analysis revealed that the PH-1 and PH-1 + DON supplementation floral infections had significantly greater disease progression than the *ΔTri5*, *ΔTri5* + DON, DON only, and mock-inoculated treatments (Kruskal-Wallis, p=2.8e-^10^; Fig. 2 (b)). To quantify the levels of DON present in all treatments at the end of disease progression (day 14), a DON-ELISA test was carried out to determine the final 15-ADON concentrations. PH-1 and PH-1 + DON samples had an average DON concentration of over 30 ppm, whilst all other treatments had no detectable (<0.5ppm) DON (Fig. 2 (c)). This indicates that the addition of DON to WT inoculum did not stimulate further DON production and confirms that the PH-1 *ΔTri5* mutant is impaired in DON biosynthesis. Of note, the lack of detection of DON in the *ΔTri5* + DON and DON alone samples is likely due to the detoxification of DON by wheat plants to DON-3- glucoside, the latter is undetectable by the competitive enzyme-labelled immunoassay kit used in this study. The conjugation of DON to DON-3-glucoside, catalysed by a UDP-glucosyltransferase, *in planta* is difficult to detect through its increased molecule polarity and is thus known as a ‘masked mycotoxin’ (Berthiller et al., 2005). A visual representation of disease progression occurring in each treatment is shown in Fig. 2 (d).

**Figure 2.**
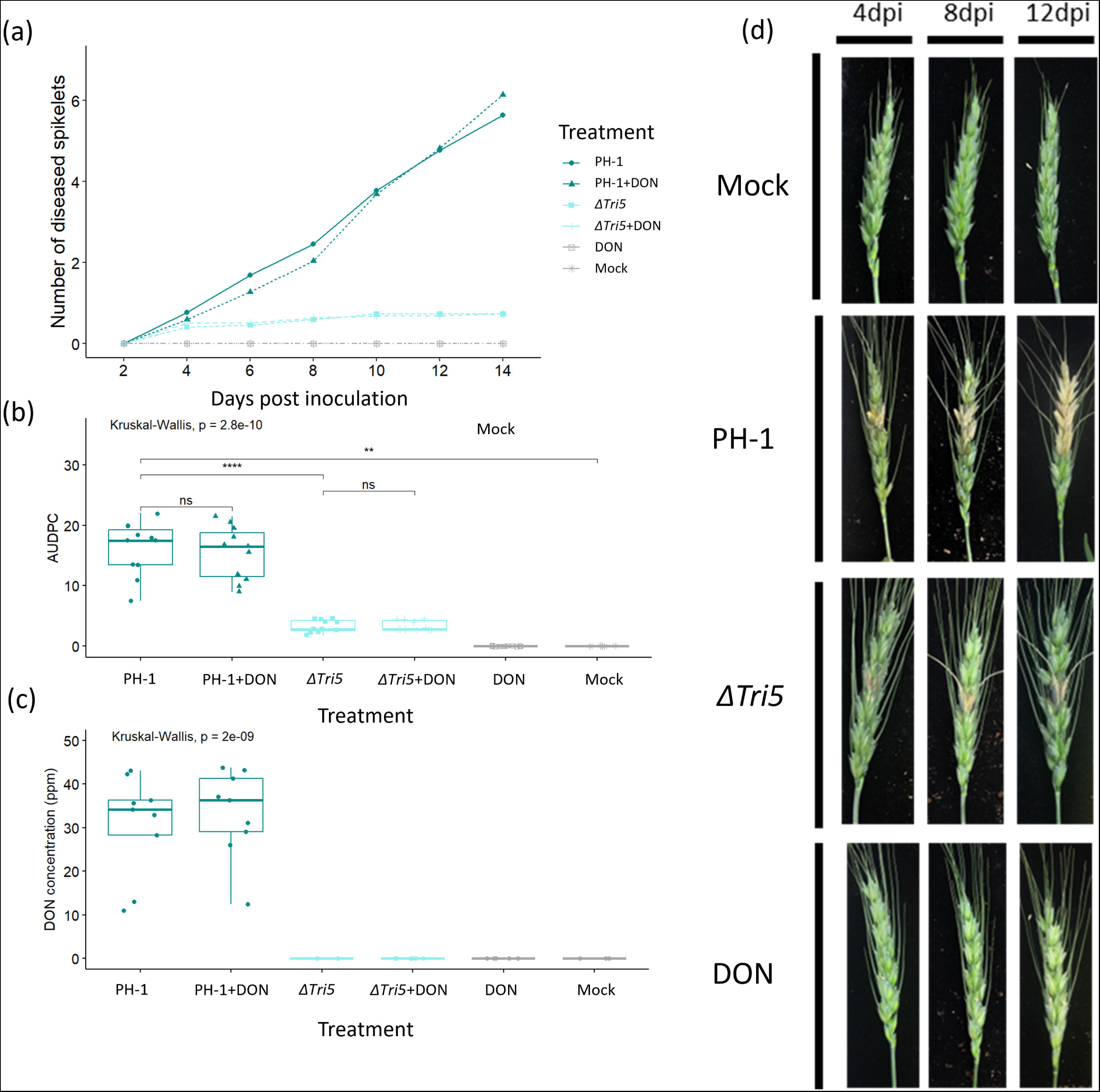
Analysis of whole wheat floral tissues following point inoculations. (a) Tracked visible disease progression at 2-day intervals to 14 dpi from below the inoculated spikelet. (b) Area Under Disease Progression Curve (AUDPC) for disease progression in panel (a), Kruskal-Wallis *p* <0.005 (***). (c) DON concentrations of wheat spikes at 14 dpi, Kruskal-Wallis *p* <0.005 (***). (d) Representative disease progression images at selected timepoints of 4, 8 and 12 days.

**Figure 3.**
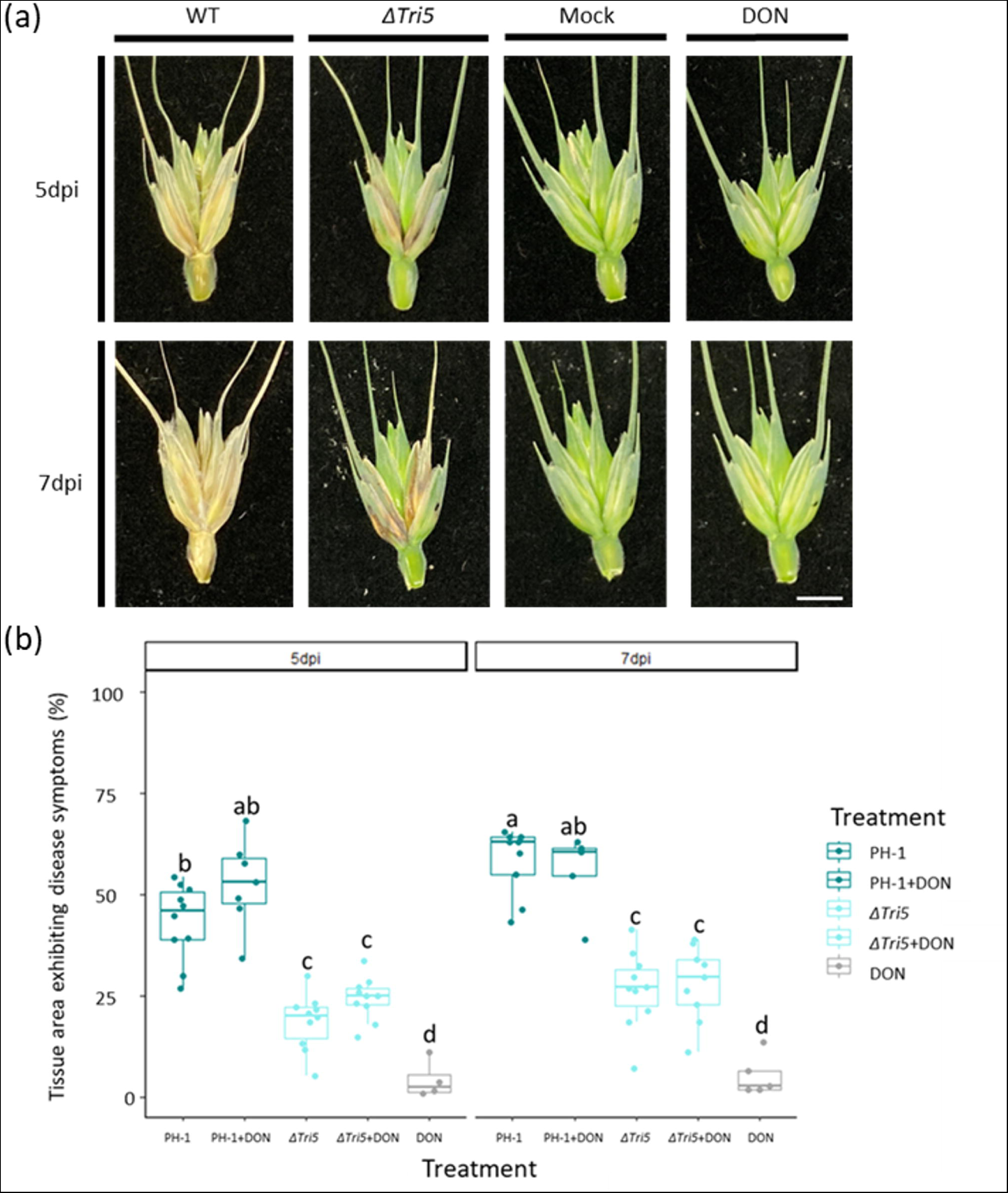
Quantitative spikelet analysis for disease symptom development. (a) Examples of dissected spikelets at 5 and 7dpi, scale bar = 10mm. (b) External tissue areas exhibiting disease symptoms at 5 and 7dpi as determined by Lemnagrid computational software. ANOVA, P < 0.005 (***), Tukey post-hoc denotes group significance.

Cuzick, Urban and Hammond-Kosack (2008) had previously shown that a qualitative difference in the appearance of macroscopic disease symptoms on the glumes between the WT and the *ΔTri5* mutant. In this study, we have extended this observation and explored the macroscopic as well as the microscopic disease symptoms. Macroscopically, we were able to confirm the *ΔTri5*-inoculated spikelets exhibited ‘eye-shaped’ lesions on the outer surface of the glume by 7dpi (Fig. 3 (a)). These differed from the characteristic fawn brown ‘bleaching’ of the spikelet tissues observed in the WT interaction at 7dpi (Fig. 3 (a)). Chemical complementation of *ΔTri5* did not restore the WT phenotype nor visibly increase the severity of the WT disease phenotype. To quantify the diseased area, inoculated spikelets were imaged at 5 and 7dpi and analysed using the Lemnagrid software for diseased area. The PH-1 and PH-1 + DON spikelets had a greater area exhibiting disease symptoms than both the *ΔTri5* and *ΔTri5* + DON treatments (Fig. 3 (b)). N. B. Computational restrictions in spikelet parsing from background led to minor, insignificant disease symptoms for DON and mock samples.

### The *ΔTri5* mutant is inhibited in its ability to traverse plasmodesmata during host-tissue colonisation

Resin-embedded samples of the lemma, palea and rachis spikelet components revealed changes in cellular morphology at different points of infection (Fig. 4). In the palea and lemma parenchyma tissue layer, the *ΔTri5* and *ΔTri5* + DON infected samples exhibited extensive cell wall degradation and colonisation by invasive hyphae (Fig. 5), similarly to the WT infection. However, in the adaxial layer of the palea and lemma tissues, the hyphae in the *ΔTri5* and *ΔTri5* + DON samples rarely penetrated into the thicker-walled cells (Fig. 5(a)-(d)). Mirroring the macroscopic lack of symptoms in the rachis, the *ΔTri5* rachis samples never contained invasive hyphae at either 5 or 7 dpi (Fig. 5(e)). In the PH-1 and PH-1 + DON infected samples, invasive hyphae proliferated throughout the entirety of the lemma, palea and rachis tissues, causing extensive cell wall degradation (Fig. 4). To penetrate the adaxial layer, the PH-1 hyphae utilised cell wall pits resembling PD pit fields (Fig. 4(c)). In these instances, the hyphae constricted considerably to traverse the cell wall. Traversing of the cell wall through PD pit fields was not observed in *ΔTri5* and *ΔTri5* + DON samples at either time point (Fig. 5). In general, where hyphae had invaded cells, the cell contents, notably nuclei, chloroplasts and evidence of cytoplasm, were not observed, indicating cell death. In the palea and lemma tissues of the PH-1 infected samples at 7dpi, putative evidence of ‘ghost’ hyphae was identified, which are characterised by a lack of cellular contents (Brown et al., 2011) in older infection structures as the infection front advances into the host plant.

**Figure 4.**
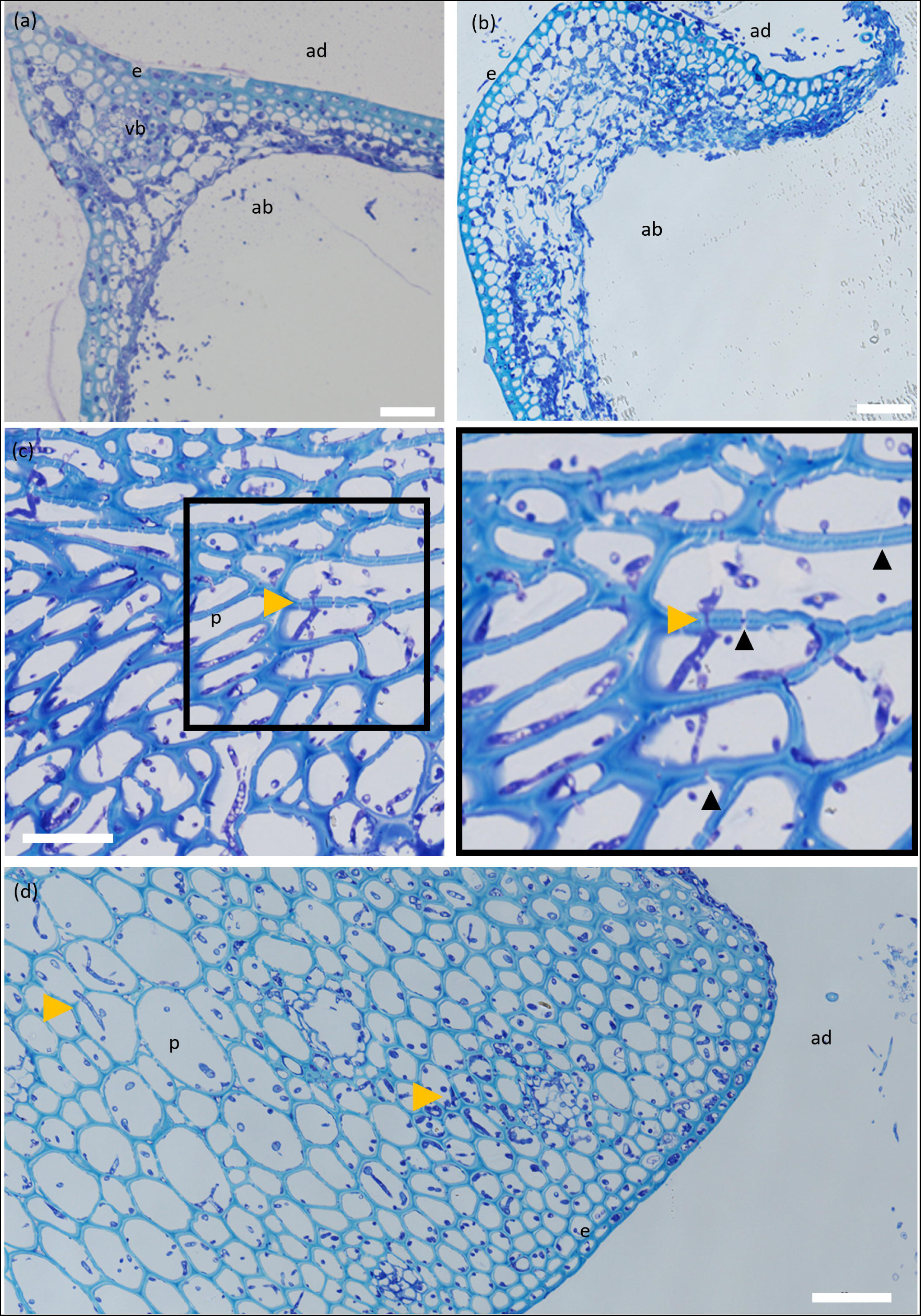
WT-infected wheat floral tissues at 5 and 7 dpi demonstrating aspects of typical infection. (a) Lemma at 5dpi infected with WT *F. graminearum* showing widespread hyphal colonisation throughout the tissue accompanied by hyphal proliferation protruding from the abaxial layer. (b) A 7dpi WT-infected lemma showing further tissue degradation by cell wall degrading enzymes and considerable hyphal proliferation. (c) Rachis at 5dpi infected with WT *F. graminearum* showing a number of plasmodesmatal crossings by invasive hyphae, indicated by yellow arrowheads, and extensive cell wall degradation of the mesophyll layer by *F. graminearum-*secreted cell wall degrading enzymes. Plasmodesmata can be identified as gaps in the parenchyma layer cell walls, a number of which are indicated by black arrowheads. (d) A 7dpi WT-infected rachis demonstrating durability of parenchyma tissue against cell wall degrading enzymes at later infection timepoints. ab = abaxial layer, ad = adaxial layer, e = epidermal layer, mes = mesophyll, p = parenchyma tissue, vb = vascular bundle. Yellow arrowheads indicate plasmodesmatal crossings by invasive *F. graminearum* hyphae. Scale bar = 50µm.

**Figure 5.**
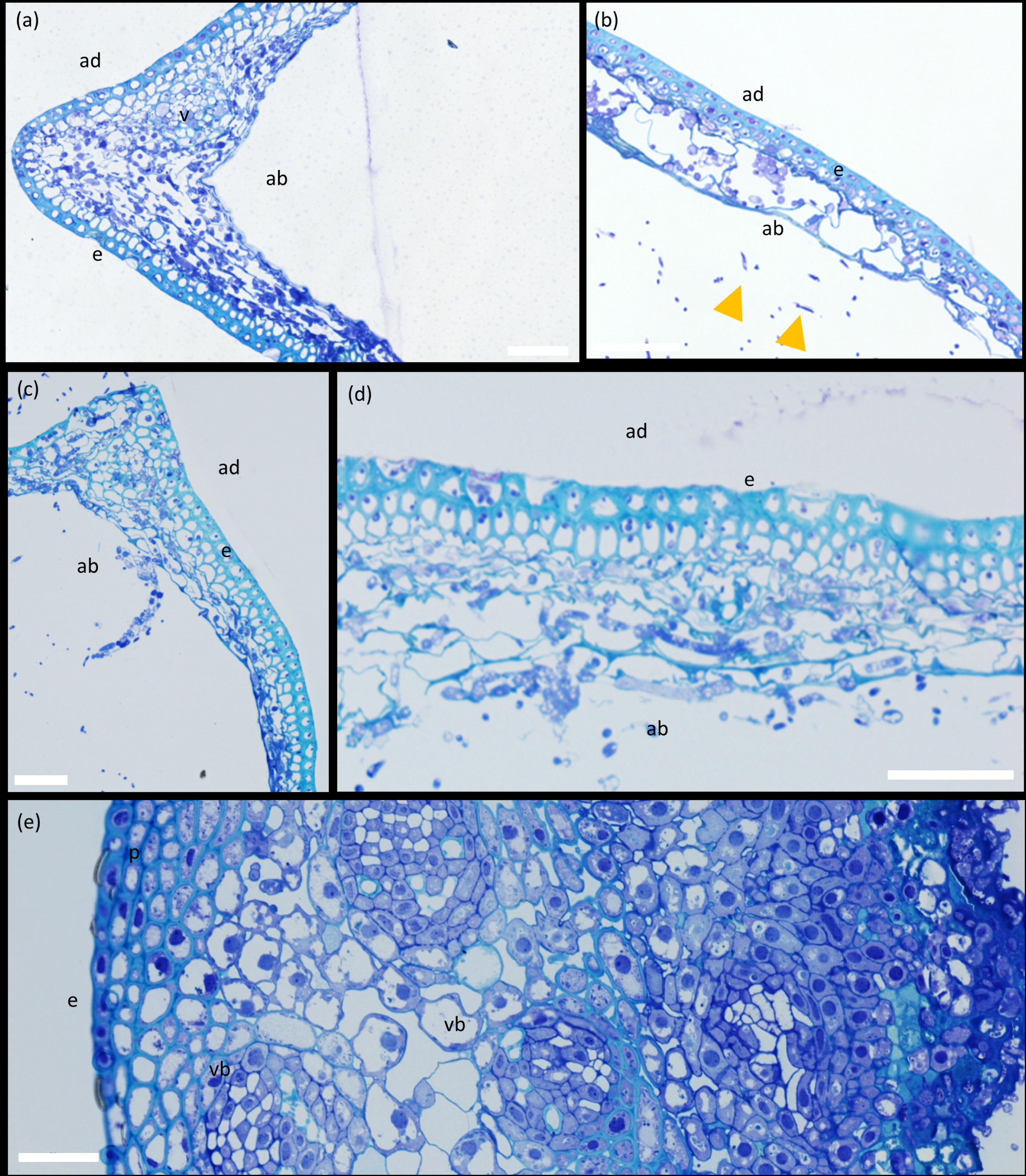
Comparison of *ΔTri5*-infected and *ΔTri5* + DON infected wheat floral tissues at 5dpi and 7dpi showing the similarities and differences between tissue types in various aspects of a typical infection. (a) Lemma at 7dpi infected with *ΔTri5 F. graminearum* with extensive proliferation of invasive hyphae throughout the abaxial layer, but rarely any penetration into the adaxial layer. (b). Palea at 5dpi infected with *ΔTri5* and supplemented with 35ppm DON showing cell wall degradation in the abaxial layer and evidence of external fungal hyphae. Yellow arrows indicate hyphae external to the plant tissue. (c) Palea at 7dpi infected with *ΔTri5*, with similar symptoms to the lemma at the earlier 5dpi time point. (d) Lemma infected with *ΔTri5* and supplemented with 35ppm DON at 7dpi showing cell wall degradation of the abaxial layer. (e) A rachis section at 5dpi infected with *ΔTri5* and supplemented with 35ppm DON. No evidence of hyphae or cell wall degradation throughout the sample. ab = abaxial layer, ad = adaxial layer, e = epidermal layer, mes = mesophyll layer, p = parenchyma tissue, vb = vascular bundle. No plasmodesmatal crossings by invasive *F. graminearum* hyphae are evident. Scale bar = 50µm.

To aid elucidation of the role of DON during infection of wheat floral tissues, cell wall thickness from resin-embedded wheat samples was measured along the adaxial layer of lemma and palea tissues, and in the visibly reinforced regions of rachis tissue, for all treatments. In the adaxial layer of the lemma, palea and rachis tissues, cell wall thickness was found not to differ between treatments, particularly between those with and without the presence of DON (Supplementary file S5). This unanticipated result indicates that cell wall reinforcements are not evident at this level of resolution, and are not impacted by the presence of DON. However, it is worth noting that extensive cell wall degradation was present in the abaxial layer of palea and lemma tissues. This microscopic phenotype was not quantified but is most likely caused by the release of CWDEs from *F. graminearum* hyphae (Supplementary file S5).

In order to gain a thorough understanding of infection, a scanning electron microscopy analysis was used. SEM micrographs of rachis post spikelet inoculation with WT PH-1 at 5dpi revealed several notable interactions, including intracellular growth through cells still containing cytoplasm, apoplastic growth between cells, hyphal constriction and cell wall traversing and gaps in rachis cell walls (Fig. 6(a), (b) and (d)). Micrographs of *ΔTri5-*infected lemma tissue at 5dpi confirmed the resin analysis, whereby extensive cell wall degradation was observed in the parenchyma tissue layer (Fig. 6(c)).

**Fig. 6.**
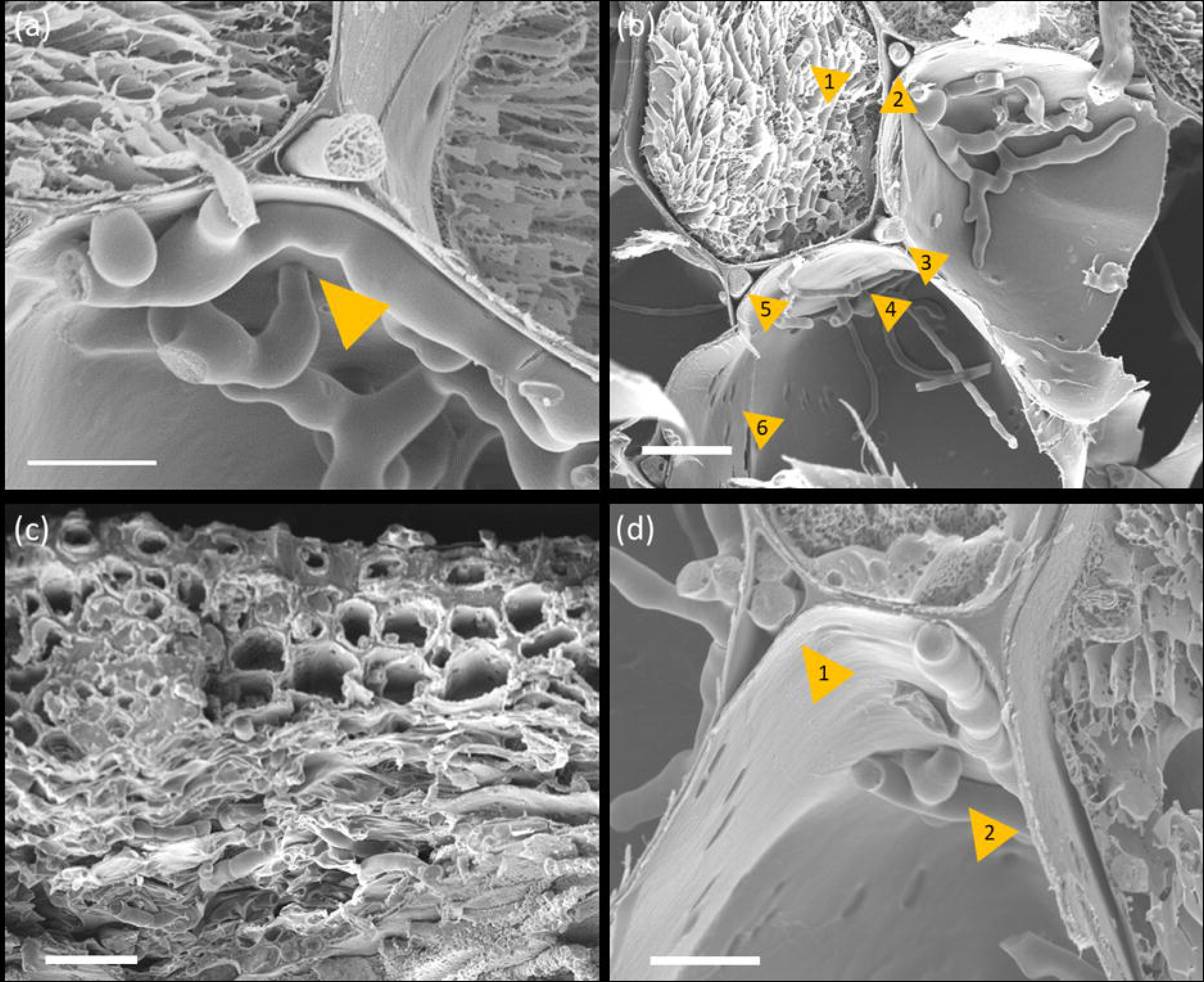
SEM micrographs of PH-1 and *ΔTri5*- wheat floral interactions. (a) A hypha of the wild-type PH-1 strain appears to cross through the cell wall at 5dpi in rachis tissue. Scale bar = 10µm. (b) Wild-type PH-1 infecting rachis tissue at 5dpi, the numbered yellow arrowhead indicates point of interest. 1. Intracellular growth in a cell where cytoplasm is still present; 2., 3., and 5. Apoplastic growth between cells, 4. Potential crossing of the cell wall by a hypha through a plasmodesma, and 6. ‘Holes’ in the cell wall that are potential sites of plasmodesmata. Scale bar = 20µm. (c). *ΔTri5-*infected lemma tissue at 5dpi demonstrating extensive hyphal colonisation and cell-wall degradation of the parenchyma tissue layer (bottom), but minimal infection in the thicker-walled adaxial layer (top), scale bar = 20µm. (d) Wild-type PH-1 infection of the rachis at 5dpi, 1. Growth of two hyphae through the same apoplastic space in parallel to hyphae growing intracellularly in neighbouring cells to the left and right. 2. Hypha appear to constrict to traverse the cell wall. Scale bar = 10µm.

### Immuno-labelling of callose during infection reveals reduced deposits in the WT infection and phloroglucinol staining indicates lignin-based defence response(s)

Resin sections of PH-1, *ΔTri5* and mock-inoculated wheat floral tissues were analysed for the presence of callose at junctions in the cell wall (Fig. 7). Immuno-labelling for the presence of callose confirmed the material of pit structures was consistent with plasmodesmata. Imaging revealed that in both WT (PH-1) and DON-deficient (*ΔTri5*) *F. graminearum*-inoculated spikes there was an increased frequency of instances where callose was deposited at plasmodesmatal junctions compared to mock-inoculated controls (Fig. 7). However, the DON only inoculated samples exhibited a marked increase in callose in both lemma and rachis tissues, indicating that callose deposition had been induced in a manner consistent with a basal immune response to symplastically isolate cells after the detection of the DON toxin.

**Fig. 7.**
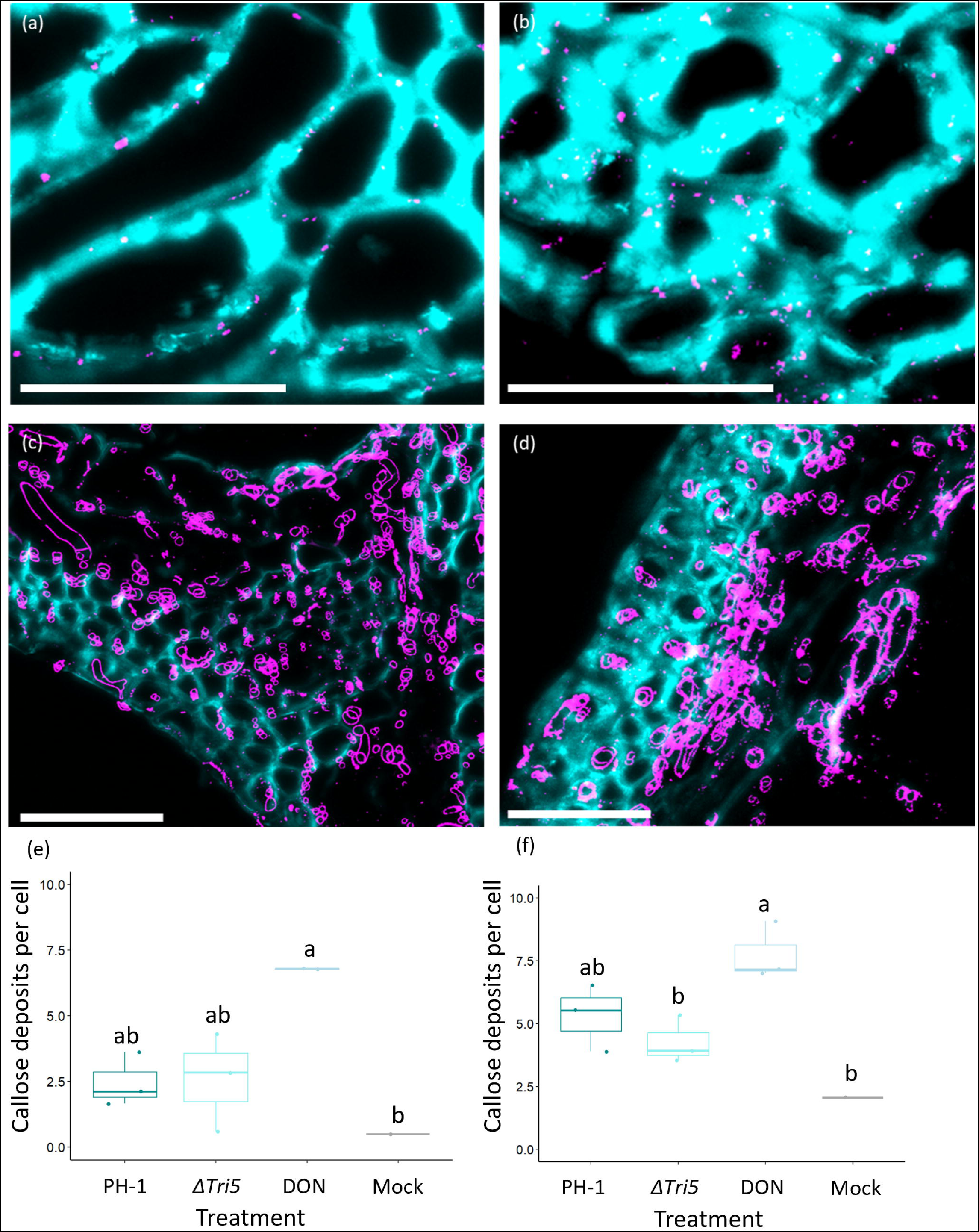
Immunofluorescence detection of callose in rachis and lemma tissues. Magnified region of interest of the *Fg-*wheat interaction demonstrating callose deposits at plasmodesmata. (a) Control rachis, (b) rachis below DON-inoculated spikelet, (c) PH-1 infected lemma at 5dpi, (d) *ΔTri5*-infected lemma at 5dpi. Sections were imaged by confocal microscopy with excitation-emission spectra for AlexFluor-488 488nm, 510nm-530nm and 405nm, 450nm-475nm for Calcofluor. Callose deposits are labelled in magenta with wheat cell walls in cyan. Scale bars = 50µm. In panels (c) and (d) the Fusarium hyphae also react positively to the antibody due to ꞵ-1,3-glucans in the fungal cell wall and are labelled in magenta. (e) Quantification of the number of immuno-labelled callose deposits, averaged across number of cells in the sample area, in lemma tissues at 5dpi, ANOVA = p < 0.05 (*), and (f) Rachis tissues at 5dpi, ANOVA = p < 0.05 (*). Letters indicate significance differences between groups from Tukey Post-hoc analysis following one-way ANOVA.

Spikelets of wheat inoculated with WT, PH-1 and *ΔTri5* were sampled at 5dpi for analysis of the lignin response. This investigation was prompted by the presence of localised ‘eye-shaped’ lesions in the *ΔTri5-*infected samples. Darker staining by the phloroglucinol indicates a higher lignin content, which was found to be most notable in the *ΔTri5-*infected lemma tissue (Supplementary file S8). This was surprising, as the lesions were present on the glume. Whilst this was not quantified, the WT PH-1 and mock-inoculated controls were visually comparable, indicating that WT *F. graminearum* may have a role in dampening pathogen-induced lignin upregulations, possibly through the action of DON. This proposes the hypotheses that in the absence of trichothecene mycotoxins, wheat is able to upregulate lignin defence pathways.

## Discussion

This study has re-examined and extended knowledge on the restricted host tissue colonisation phenotype previously reported in wheat spikes for the non-DON-producing *ΔTri5* single gene deletion mutant of *F. graminearum*. The study was catalysed by the lack of published cellular information available on how DON, produced and secreted by the advancing *F. graminearum* hyphae, actually facilitates the extraordinary effective and speedy disease progression consistently observed in the spikes of susceptible wheat cultivars. DON has long been classified as a key virulence factor in the *F. graminearum*-wheat interaction (Hohn et al., 1993; Jansen et al., 2005) and facilitates the host-tissue colonisation of the rachis and thus is essential for successful internal spikelet-to-spikelet growth of hyphae through the entire floral spike. However, prior to this study, the morphological and cellular responses underlying this macroscopically well documented phenomenon had not been explored. In this study our two primary aims were (a) to identify the morphological differences in the hyphal infection routes between the wild-type (WT) and *ΔTri5* strains during wheat floral infections compared to coleoptile infections, and (b) to focus on the multifaceted role(s) of the cell wall and its constituent components as well as the pit fields during hyphal colonisation due to their potential to delay, minimise or cease fungal progression through the numerous internal complexities that the wheat spike architecture presents to the *Fusarium* hyphae.

As described above, our experimentation confirmed that the *ΔTri5* mutant could sufficiently colonise the lemma and palea tissues but not the rachis (Jansen et al., 2005; Cuzick et al., 2007). Similarly, our results concurred with those of Jansen et al. (2005) that the DON-deficient *F. graminearum* strain could not grow beyond the rachis node due to the presence of inherently thicker cell walls in this tissue. However, our quantitative comparative analysis of the WT and DON-deficient interactions revealed no differences between cell wall thickness at two timepoints, or with the control mock inoculated tissues, indicating that cell walls do not increase in thickness *per se* as part of a locally occurring defence response. Upon further microscopic analysis in the current study, we observed that the DON-deficient *ΔTri5* mutant could not enter wheat cells with inherently thicker cell walls because the hyphae could not pass through pit fields containing plasmodesmata. This phenomenon was frequently observed in both the cortical and sclerenchyma cell layers. As a result, the *ΔTri5* hyphae accumulated within and between the neighbouring thinner-walled parenchyma cells. In the absence of DON, potentially other so far uncharacterised secreted proteinaceous effectors fail to correctly manipulate these potential gateways into the neighbouring wheat cells. The analysis of resin sections revealed that cell walls within the adaxial layer of lemma and palea tissues were not thicker in infected samples. Although this rules out additional cell wall reinforcements, these findings do not eliminate cell wall compositional changes. Our results indicate that lignin content increases in the lemma tissue, which strengthens the tissue and hence emphasises the role of plasmodesmata as cell wall portals in host-tissue colonisation. We hypothesise that DON, through its intracellular target of the ribosomes, inhibits local protein-translation based defence responses.. Whereas symplastic isolation of neighbouring cells by the deposition of callose at plasmodesmata is a largely post-translationally regulated process induced within the generic plant defence response (Wu et al., 2018). Our SEM inquiry of the infected tissues indicates that plasmodesmata, when used by the advancing hyphal front, are potentially ‘dead portals’, that lack the desmotubule symplastic bridge between neighbouring cells. Whether callose deposits are eliminated prior to or coincident with hyphal constriction and traversing of the cell wall will require the use of another microscopy technique, namely TEM. Although again a static analysis method, TEM can be used to explore whether desmotubule connections, and callose deposits, are consistently present or absent at the point of hyphal traverse. Collectively, these data suggest that the broad-spectrum consequences of DON targeting could prevent the synthesis and action of key defence enzymes at plasmodesmata. This could be explored by a combined comparative proteomics, phosphoproteomics and RNAseq analysis of the WT and *ΔTri5*-infections to elucidate the wheat defence responses occurring at the advancing *Fusarium* hyphal front that are reduced and/or eliminated by the presence of DON.

The deposition of callose at the plasmodesmatal junction by callose synthases has been demonstrated to be induced by various biotic stress-inducing pathogens (Wu et al., 2018). The role of callose differs with cellular location: callose polymers are a structural component of papillae in various cereal species that form below appressoria produced by fungal pathogens such as the powdery mildew *Blumeria graminis* f. sp*. hordei*, whereby elevated callose deposition in highly localised papillae in epidermal cells result in resistance to fungal infection (Ellinger et al., 2013). In vascular tissue, callose can be deposited to restrict vascular advancements by wilt pathogens, including by *Fusarium* and *Verticillium* species (reviewed: Kashyap et al., 2021). To investigate the potential of DON impacting upon plasmodesmatal occlusion following our discovery of the impeded traversal of plasmodesmata by the *ΔTri5* strain, we immuno-labelled callose in resin-embedded sections of wheat floral tissues. We found that DON strongly induced callose depositions, and callose deposition was also moderately increased in WT and *ΔTri5* infected lemma and palea tissues. This indicates that callose deposition is increased as a defence response when DON or *Fusarium* hyphae are present. However, in the WT infection, we observed a frequency of callose depositions similar to the non-DON producing *ΔTri5* strain indicating an interruption or targeted degradation of callose occlusions by *F. graminearum* invasive hyphae. The secretion of glycoside hydrolase (GH) proteins that break down ꞵ-1,3-glucans such as callose have not been explored with respect to the *Fg*-wheat interaction, although GH12 family proteins that break down xyloglucan in plant cell walls appear to be implicated in virulence (Wang et al., 2022). In the *ΔTri5* infections, in the absence of DON other hyphal components and /or secreted molecules may be responsible for the modest callose deposition at the plasmodesmatal junction.

Intracellular colonisation through the rachis node and beyond in the rachis internode possibly requires DON and is therefore required for the second intracellular phase of the biphasic lifestyle described for *F. graminearum,* where extracellular apoplastic growth characterises the initial ‘stealth’ phase of infection (Brown et al., 2010). If this is the case, then lacking the ability to traverse plasmodesmata would restrict direct acquisition of nutrients from host cells by the fungal hyphae. The *TRI* biosynthetic gene cluster required for DON biosynthesis is transcriptionally activated early during wheat spike infection, peaking between 72 and 120 hrs post-inoculation (Evans, 2000; Cheng, Kistler and Ma, 2019), when infection is largely restricted to the palea, lemma and glume tissues. Trichothecene biosynthesis is regulated by two transcription factors, TRI6 and TRI10, within the biosynthetic pathway. Of note, DON is not required for full virulence of the developing wheat kernel seed coat (Jansen et al., 2005), in addition to our finding in coleoptiles, Ilgen et al. (2009) identified that trichothecene biosynthesis pathway induction was potentially tissue specific and somewhat restricted to the developing grain kernel and rachis node, and suggested that ‘kernel tissue perception’ by the *Fusarium* hyphae induce the biosynthesis of trichothecene mycotoxins. This suggestion concurs with Zhang et al. (2012) who report that the trichothecene biosynthesis genes were not induced during *F. graminearum* infection of wheat coleoptiles, whereas Qui et al. (2019) reported the expression of *TRI4* in wheat coleoptiles. We found *TRI5* expression to be detectable at low levels in the wheat coleoptiles, remaining around 0.8 x the expression levels of FgActin. This is lower than the TRI5 expression level reported by Brown et al. (2011), who reported a level over 3 x that of FgActin in wheat spikes.

Gardiner et al. (2009b) presented evidence that, in addition to their previous reports that exogenous application of amines, such as agmatine, *in vitro* induces *TRI5* expression (Gardiner et al., 2009a), low pH further accelerates expression of the *TRI* gene cluster. Other inducers of the DON biosynthetic pathway genes include carbon, nitrogen and light (Gardiner et al., 2009b). These factors could explain the low and stationary levels of *TRI5* gene expression throughout infection of wheat coleoptiles, which have been noted to be particularly acidic plant tissue due to the optimum activity of expansion proteins around pH 4 (Gao et al., 2008). Other fungal pathogens that are reported to utilise plasmodesmata during infection of cereals include *Magnaporthe orzyae* and *M. oryzae* pathotype *triticum* which respectively cause rice blast and wheat blast diseases on the floral panicles and floral spikes (Sakulukoo et al., 2018, Fernandez and Orth, 2018). Although *Magnaporthe oryzae* does not synthesise trichothecene mycotoxins, the invading hyphae secrete another potent general protein translation inhibitor, namely tenuazonic acid (Wilson and Talbot, 2009). The effect of this mycotoxin on plasmodesmatal traversing and virulence in *Magnaporthe spp*. has not yet been reported.

*F. graminearum* progression into the rachis and through sequential rachis nodes and internodes allows for the successful completion of the disease infection cycle in wheat crops. Typically, perithecia form from the chlorenchyma band of the rachis following prolific hyphal colonisation of this highly specialised photosynthetic tissue layer within the wheat spike (Guenther and Trail, 2005). Hence, interruption of WT disease progression prior to this crucial point in the primarily monocyclic infection cycle is of great interest for reducing full virulence of FHB and in particular in reducing the abundance of air dispersed ascospores. Interestingly, infection of barley spikelets with WT *F. graminearum* is solely restricted to the inoculated spikelet, similarly to *ΔTri5* infection of wheat (Jansen et al., 2005). How this occurs has not yet been explored, but we hypothesise that a lack of traversing of plasmodesmata by hyphae may have a role to play in barley rachis node tissue.

Overall, our study indicates that plasmodesmata are the key to successful host-tissue colonisation by *F. graminearum* and that DON, directly or indirectly, facilitates this interaction. We anticipate that the results of this study are considered in future working disease models of the *F. graminearum* -wheat interaction and suggest these incorporate a greater emphasis on tissue and cell wall architecture and composition when considering host susceptibility to fungal pathogens. To this end, we have proposed a new working model (Fig. 8), that summarises our findings around the presence of DON during wheat infection and the impact on callose deposition at plasmodesmata.

**Fig. 8.**
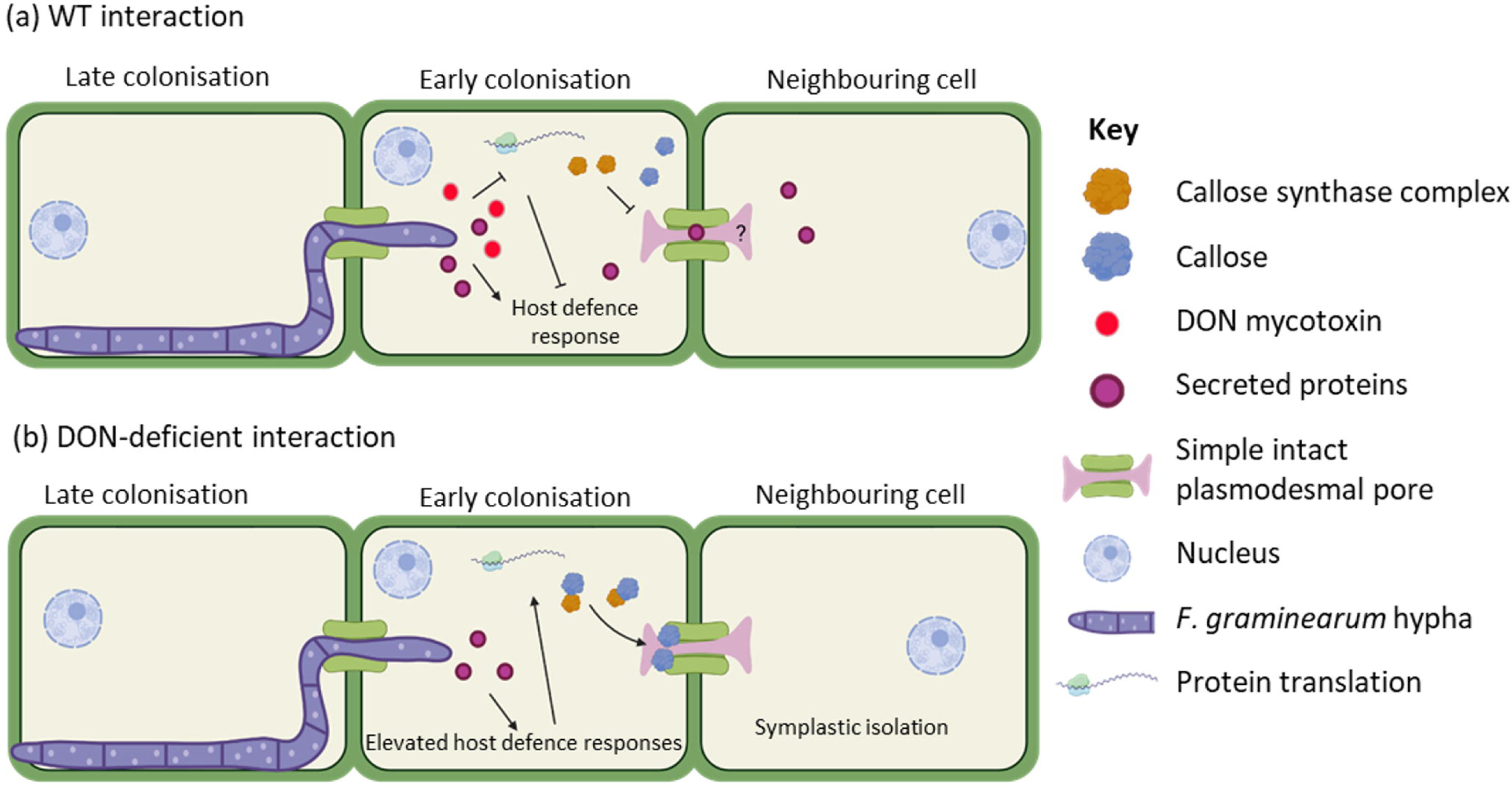
Proposed working model for the role of DON in the *F. graminearum* – wheat interaction. (a) In the wild-type (WT) interaction, DON interferes with the wheat host defence response by inhibiting protein translation and reducing the ability of the host to deposit callose at plasmodesmata to restrict further hyphal infection. It is currently unknown how long the desmotubule remains functional, or in place. (b) In the absence of DON, *Fg*-secreted proteins are detected by the host and trigger host defence responses, including the symplastic isolation of neighbouring cells by the deposition of callose at plasmodesmata.

## Experimental Procedures

### Fungal growth

The *Fusarium graminearum* reference strain PH-1 (NCBI: txid229533) and the DON-deficient single gene deletion mutant *ΔTri5*, within the PH-1 parental background (Cuzick et al., 2008), were used in this study. Conidia for glycerol stocks were prepared by culturing on Synthetic Nutrient Poor Agar (SNA) plates containing 0.1% KH_2_PO_4_, 0.1% KNO_3_, 0.1% MgSO_4_ · 7H_2_O, 0.05% KCl, 0.02% glucose, 0.02% sucrose and 2% agar. Plates were left to grow for 8 days at room temperature (RT) with constant illumination under near-UV light (Philips TLD 36W/08). TB3 liquid medium (0.3% yeast extract, 0.3% Bacto Peptone and 20% sucrose) was added to plates to stimulate spore production and left for a further 2 days. Conidia were harvested and stored in 15% glycerol at −80°C in 2ml cryotubes (Thermo Fisher Scientific, MA, USA). Conidial suspensions in water to be used for inoculations were prepared by spreading conidia from glycerol stocks onto Potato Dextrose Agar (PDA, Sigma Aldrich, UK) plates, then growth at RT for 2 days, harvesting with dH_2_O and spore concentrations measured with the aid of a haemocytometer (Hausser Bright-line, USA). Experiments were conducted under APHA plant licence number 101948/198285/6.

### Plant growth

The susceptible dwarf spring wheat (*Triticum aestivum*) cultivar (cv.) Apogee was used for all wheat experiments, sourced from the National Small Grains Collection, USDA-ARS, Aberdeen, Idaho, USA. Seeds were sown in Rothamsted Prescription Mix (RPM) soil (Petersfield Growing Mediums, UK) in P15 pots (approx. volume 7cm^3^) and grown in controlled environment facilities at HSE category 2 (Fitotron®, Weiss Gallenkamp, UK), 16hr light: 8hr dark cycle at 22°C and 18°C respectively, 70% relative humidity and illumination at 2.2×10^3^ µmol m^-3^.

### Coleoptile inoculations

For coleoptile inoculations, Apogee grain were left for 2 days at 5°C in water for imbibition before being placed individually onto cotton wool plugs in a 24-well tissue culture plate (VWR, USA) and left to germinate for 3 days under high humidity conditions (<90% relative humidity) under normal wheat growth conditions. At 3 days post-sowing, approximately 5mm from the tip of each coleoptile was cut to encourage infection. Inoculations occurred through the placement of a cut pipette tip with a filter paper insert soaked with 5×10^5^ spores/ml solution, with dH_2_O used as a negative control. The coleoptiles were left in the dark for 3 days to aid infection, after which inoculation tips were subsequently removed, and coleoptiles were left to grow under normal growth conditions for a further 4 days (Darino et al., 2024). Disease phenotypes on the coleoptiles were assessed at 7 days post inoculation by imaging lesions on a Leica M205 FA Stereomicroscope (Leica Microsystems, UK). Each experimental replicate contained 5 biological samples for each treatment (3 mock-inoculated) and the experiment was repeated 3 times.

### Floral inoculations

At mid-anthesis, wheat plants were inoculated with 5 ×10^5^ spores/ml water conidial suspension of PH-1 or *ΔTri5*, conidial suspension supplemented with DON, DON alone or water (dH_2_O) control. DON supplementation of inoculum was 35ppm (Sigma-Aldrich, USA). As described in Lemmens et al. (2005), a 5µl droplet was placed between the palea and the lemma on each side of the 7^th^ true spikelet from the base. Inoculated plants were placed in a high (<90%) humidity for the first 72 hours of infection, with the first 24 hours in darkness. After 72 hours plants were returned to normal growth conditions.

### Determination of DON concentration for exogenous application

To determine the concentration of DON to utilise for the chemical complementation experiments, a number of experiments were undertaken. These scoped for a concentration that was not detrimental to both *F. graminearum* spore germination and growth (supplementary S1), as well as sufficient to cause physiological stress to plant tissue (supplementary S2). Concentrations up to 350ppm were non-detrimental to fungal proliferation in media and as low as 10ppm demonstrated considerable reduction in growth in *Arabidopsis thaliana* seedlings. The concentration of 35ppm (118μM) was determined as a result of these experiments.

### Disease progression

As above, Apogee at mid-anthesis was inoculated and disease progression was assessed by counting spikelets showing visible symptoms every 2 days after inoculation until 14 dpi. Area Under Disease Progression Curve (AUDPC) (Van der Plank, 1963) values were calculated using the ‘agricolae’ package (version 1.4.0, de Mendiburu and Yaseen, 2020) in R (version 4.0.2). Statistical significance was determined by Kruskal-Wallis one-way analysis of variance through the R package ‘ggplot2’ (version 3.4.0, Wickham, 2016).

### Red Green Blue (RGB) colour classification for disease assessment of dissected spikelets

To quantify disease progression on wheat spikelets at 5 and 7dpi, colour (RGB) spikelets were imaged (iPhone 6s, Apple Inc, US) on both sides with consistent illumination. Diseased area was quantified using a curated programme on the LemnaTec Lemnagrid software (CHAP, York, UK). Diseased area was classified by pixel colour segmentation after application of filters to threshold from the background, identify misclassified pixels and fill in gaps. Area attributed to anthers were omitted from further analysis. The relative area attributed to each classification was then calculated in a custom R script and all samples were normalised to the mean value of ‘diseased’ of the mock treatment due to background parsing error.

### Bioimaging

Inoculated spikelets were dissected from the wheat spikes for internal observations of infected floral tissues. Spikelets were fixed for 24 hr in a solution of 4% paraformaldehyde, 2.5% glutaraldehyde and 0.05M Sorensen’s phosphate buffer (NaH_2_PO_4_: Na_2_HPO_4_ · 7H_2_O, pH 7.2), in the presence of Tween 20 (Polyethylene glycol sorbitan monolaurate; Sigma-Aldrich) and subject to a light vacuum for 20s to ensure tissue infiltration. Fixed spikelets were washed 3x with 0.05M Sorensen’s phosphate buffer and subsequently underwent an ethanol dehydration protocol at 10% EtOH increments, up to 100% EtOH. Spikelets were dissected into component tissues and embedded with LR White resin (TAAB, Reading, UK) at increasing resin ratios (1:4, 2:3, 3:2, 4:1), followed by polymerisation in the presence of N_2_ at 60°C for 16h. Ultra-thin 1µm resin sections were cut from resin blocks using a microtome (Reichert-Jung, Ultracut), placed onto Polysine microscope slides (Agar Scientific, UK) and stained with 0.1% (w/v) Toluidine Blue O in 0.1M Sorensen’s phosphate buffer (NaH_2_PO_4_: Na_2_HPO_4_ · 7H_2_O, pH 7.2). Every 10^th^ section was collected for a total of 10 sections per embedded block to fully explore floral tissues and mounted with Permount (Fisher Scientific, UK) prior to imaging on a Zeiss Axioimager 512 (Zeiss, Oberkochen, Germany) at x20 magnification under brightfield illumination. The experiment was repeated 3 times, with a total of 5 biological replicates for each treatment, with 2 mock samples per batch. In total, 111 resin blocks were explored across a 100um in the centre of the sample, with sections cut every 10µm. Image analysis was conducted in Fiji for ImageJ (version 2.3.0) and statistical analysis was conducted in R (version 4.0.2).

For SEM exploration of floral tissues, spikelets at 5dpi were excised and coated in 50:50 OCT compound (Sakura FineTek) with colloidal graphite (TAAB). SEM analysis was conducted on rachis tissue infected with the WT reference strain PH-1 at 5dpi. Sample preparation occurred in a Quorum Cryo low-pressure system before imaging on a JEOL LV6360 SEM at 5kV with software version 6.04.

Callose immuno-labelling of resin-embedded sectioned material was conducted according to Amsbury and Benitez-Alfonso (2022). Briefly, callose was localised by anti-ꞵ-1,3-glucan antibodies (Biosupplies, Australia) and secondarily conjugated with rabbit anti-mouse Alexa Fluor 488. Wheat cell walls were counterstained with calcofluor white. Sections were imaged by confocal microscopy on a Leica SP8 confocal microscope, with excitation-emission spectra for AlexFluor-488 at 488nm, 510nm-530nm and 405nm, 450nm-475nm for Calcofluor white. Image analysis for the quantification of callose deposits per cell was conducted in Fiji (ImageJ) using maximum projections of Z stacks and channels converted to binary masks. The number of cells in the sample area was calculated using the cell counter tool and callose deposits were counted by the number of discrete Alexa 488-fluorescences between the size of 2 to 12 pixel units to eliminate cross-reactivity with β-1,3-glucans in the fungal cell walls. The number of callose deposits were averaged across the number of complete cells (all cell walls visible) in the sample area. Callose deposits were quantified in the lemma and rachis tissues only, with 3 biological replicates for each treatment (PH-1, *ΔTri5*, DON, Mock). Further examples are present in supplementary file S6 and image analysis methodology is demonstrated in supplementary S7.

### DON quantification

To determine if the presence of DON in the WT strain inoculum stimulated further DON production, if administered DON could be detected in wheat spike tissues at the end of disease progression (14dpi), and the absence of DON in the *Δtri5* mutant interaction a competitive enzyme-labelled immunoassay for 15-ADON was employed. Whole wheat spikes were utilised for this quantification in the case the DON was trafficked or secreted beyond the inoculated spikelet due to its high water solubility. Whole wheat spikes after 14 days of disease progression were ground to a fine powder in the presence of liquid nitrogen and 1g of each sample was resuspended in 5ml dH_2_O, vortexed until dissolved, incubated in a 30°C water bath for 30 mins and centrifuged for 15 minutes at full speed (13.1g). The supernatant was removed and analysed using the Beacon Analytical Systems Inc Deoxynivalenol (DON) Plate Kit (Cat. 20-0016) according to kit instructions. The OD450 values were measured on a Thermo Varioskan microplate reader (Thermo Scientific, USA). Three technical replicates of each biological replicate (a single wheat spike) were conducted, and the experiment was repeated three times.

### Expression of the mycotoxin biosynthesis gene TRI5 during coleoptile infection

The trichodiene synthase gene *TRI5* was used as a proxy for the relative expression of the trichodiene biosynthesis pathway during coleoptile infection. Total RNA was extracted from whole coleoptiles at 3, 5 and 7dpi using a total RNA extraction kit (NEB) and following the kit instructions. First strand cDNA was synthesised using RevertAid First Strand cDNA synthesis kit (ThermoFisherSci) as per kit instructions and utilising random hexamer primers provided. *TRI5* expression was then assessed by qPCR with melt curve using the primers in supplementary S3, with SYBR as the reporter, passive reference as ROX and NFQ-MGB as the quencher. The qPCR with melt curve was conducted in technical and biological triplicate on a QuantStudio™ 6 Pro and results analysed on the complementary Design & Analysis Software v. 2.6.0 (ThermoFisher Scientific, MA, USA). The experiment was conducted 3 times. Quantification of *TRI5* expression was normalised against the housekeeping gene FgActin (primers in supplementary S3) for each sample, also included within the qPCR run in triplicate. At least one primer in a primer pair for qPCR targets was designed to span an exon-exon junction to avoid annealing to residual DNA.

### Phloroglucinol staining for presence of lignin

A 3% Phloroglucinol (Sigma Aldrich, UK) - HCl solution (Weisner stain) was prepared fresh in accordance with methods by Mitra and Loqué (2014). Inoculated wheat spikelets were sampled at 5dpi and cleared in 100% EtOH for 4 days before going through a rehydration series (75%, 50%, 25% and 0% EtOH) at 1 hour per stage. Cleared spikelets were bathed in Weisner stain for 1 hour, or until staining of the tissues becomes evidently saturated. Spikelets were then imaged (OM-D E-M10, Olympus, Japan) under constant illumination and, subsequently, dissected tissues were imaged individually.

### Formation of perithecia *in vitro*

Carrot agar was prepared using the method outlined by Cavinder et al. (2012) and supplemented with DON at 35ppm (w/v) to test for the ability of the WT strain, and the DON trichothecene-deficient deletion mutant, *ΔTri5*, to develop perithecia *in vitro*, for lifecycle completion viability (Supplementary S4). Ability of perithecia to discharge ascospores in the presence of DON was assessed using the same method as described in Cavinder et al. (2012).

### Statistical analysis

Scripts were written in R (version 4.0.2) for each experimental analysis. Unless otherwise stated, ANOVA followed by Tukey post-hoc test was conducted for parametric datasets and Kruskal-Wallis for non-parametric datasets. The significance threshold was set to P < 0.05 in all cases.

## Acknowledgements

The authors would like to thank the CHAP organisation at Rothamsted Research for access to their Lemnagrid image analysis software. Special thanks are given to Hannah Walpole and Kirstie Halsey from Rothamsted Research Bioimaging department for continued training, advice and expertise. We also thank Harold Bockelman, the curator of the National Small Grains Collection, USDA-ARS, for providing the wheat variety used in this study. Kim Hammond-Kosack and Martin Urban are supported by the Biotechnology and Biological Sciences Research Council (BBSRC) Institute Strategic Programme (ISP) Grants Designing Future Wheat (BBS/E/C/000I0250) and Delivering Sustainable Wheat (BB/X011003/1 and BBS/E/RH/230001B) and the BBSRC grants (BB/X012131/1 and BB/W007134/1). Victoria Armer is supported by the BBSRC-funded South West Biosciences Doctoral Training Partnership (BB/T008741/1).

## Author contributions

VA conducted the experiments and wrote the manuscript, MU generated the *Fusarium graminearum* mutant, TA provided oversight and training for image analysis, MJD and KHK provided project oversight, experimental design and manuscript planning, development and revisions.

## Additional Information

The authors declare no financial or non-financial competing interests.

## Data availability

The research data supporting this publication are provided within this paper. Requests for materials relating to this paper should be made to Kim Hammond-Kosack at Rothamsted Research.

## Supplementary

**S1 Determination of the DON concentration to be used in point inoculations with *F. graminearum* conidia (text)**

**S2 Toxicity of DON on plant tissue (figure) S3 Primers used in this study (5’-3’) (table)**

**S4 Perithecia formation *in vitro* does not require DON (text)**

**S5 Cell wall thickness of adaxial cell layer in resin samples (figure)**

**S6 Immunofluorescence detection of callose in sectioned floral tissues (figure)**

**S7 Quantification of immuno-labelled callose deposits in wheat spikelet resin sections (figure)**

**S8 Phloroglucinol staining of infected spikelets for the detection of lignin (figure)**

## Notes

### Competing Interest Statement

The authors have declared no competing interest.

### Summary of Updates

Following reviewer comments, additional experiments have been conducted including the expression of TRI5 during coleoptile infection at 3, 5 and 7 days post inoculation, and controls added to the supplementary demonstrating the toxicity of DON.

## References

Amsbury, S. and Benitez-Alfonso, Y. (2022). Chapter 10: Immunofluorescence detection of callose in plant tissue sections. In: Benitez-Alfonso, Y. and Heinlein, M. (eds.): *Plasmodesmata: Methods and Protocols*, Methods in Molecular Biology, vol. 2457. doi: 10.1007/978-1-0716-2132-5_10.

Armer, V.J., Deeks, M.J. and Hammond-Kosack, K. E. (2024). *Arabidopsis thaliana* detached leaf assay for the disease assessment of *Fusarium graminearum*. Protocols.io. doi: 10.17504/protocols.io.261gedmwov47/v1.

Berthiller, F., Dall’Asta, C., Schuhmacher, R., Lemmens, M., Adam, G. and Krska, R. (2005). Masked mycotoxins: determination of a deoxynivalenol glucoside in artificially and naturally contaminated wheat by liquid chromatography–tandem mass spectrometry. Journal of Agricultural and Food Chemistry. 53(9), 3421–3425. doi: 10.1021/jf047798g.

Boenisch, M.J. and Schafer, W. (2011). *Fusarium graminearum* forms mycotoxin producing infection structures on wheat. BMC Plant Biology. 11: 110. doi: 10.1186/1471-2229-11-110

Brown, D. W., Dyer, R. B., McCormick, S. P., Kendra, D. F. and Plattner, R. D. (2004). Functional demarcation of the *Fusarium* core trichothecene gene cluster. Fungal Genetics and Biology. 41(4), 454–462. doi: 10.1016/j.fgb.2003.12.002.

Brown, N. A., Antoniw, J., Hammon-Kosack, K. E. (2012). The Predicted Secretome of the Plant Pathogenic Fungus *Fusarium graminearum*: A Refined Comparative Analysis. PLoS ONE, 7(4), e33731. doi:10.1371/journal.pone.0033731.

Brown, N. A., Bass, C., Baldwin, T. K., Chen, H., Massot, F., Carion, P. W. C. et al. (2011). Characterisation of the *Fusarium*-wheat floral interaction. Journal of Pathogens. Article ID 626345. doi: 10.4061/2011/626345.scholar

Brown, N. A., Urban, M., van de Meene, A. M. L., and Hammond-Kosack, K. E. (2010). The infection biology of *Fusarium graminearum* defining the pathways of spikelet to spikelet colonisation in wheat ears. Fungal Biology. 114(7): 555–571. doi: 10.1016/j.funbio.2010.04.006.

Cavinder, B., Sikhakolli, U., Fellows, K. M. and Trail, F. (2012). Sexual development and ascospore discharge in *Fusarium graminearum*. Journal Visualized Experiments. 61, 3895. doi: 10.3791/3895.

Chen, Y., Kistler, H. C., and Ma, Z. (2019). *Fusarium graminearum* trichothecene mycotoxins: biosynthesis, regulation, and management. Annual Review Phytopathology. 57, 15–39. doi: 10.1146/annurev-phyto-082718-100318.

Cuzick, A., Urban, M. and Hammond-Kosack, K. (2008). *Fusarium graminearum* gene deletion mutants *map1* and *tri5* reveal similarities and differences in the pathogenicity requirements to cause disease on Arabidopsis and wheat floral tissue. New Phytologist. 177(4), 990–1000. doi: 10.1111/j.1469-8137.2007.02333.x

Darino, M., Urban, M., Kaur, N., Machado-Wood, A., Grimwade-Mann, M., Smith, D., et al. (2024). Identification and functional characterisation of a locus for target site integration in *Fusarium graminearum*. Fungal Biology and Biotechnology. 11: 2. doi: 10.1186/s40694-024-00171-8 .

Dilks, T., Halsey, K., De Vos, R. P., Hammond-Kosack, K. E. and Brown, N. A. (2019). Non-canonical fungal G-protein coupled receptors promote Fusarium head blight on wheat. PLoS Pathogens. 15(4): e1007666. doi: 10.1371/journal.ppat.1007666.

Ellinger, D., Naumann, M., Falter, C., Zwikowics, C., Jamrow, T., Manisseri, C., et al. (2013). Elevated early callose deposition results in complete penetration resistance to powdery mildew in Arabidopsis. Plant Physiology. 161(3): 1433–1444. doi: 10.1104/pp.112.211011

Evans, C. K., Xie, W., Dill-Macky, R. and Mirocha, C. J. (2000). Biosynthesis of deoxynivalenol in spikelets of barley inoculated with macronidia of *Fusarium graminearum*. Plant Disease. 84(6): 654–660. doi: 10.1094/PDIS.2000.84.6.654.

Fan, J., Urban, M., Parker, J. E., Brewer, H. C., Kelly, S. L., Hammond-Kosack, K. E. et al. (2013). Characterization of the sterol 14α-demethylases of *Fusarium graminearum* identifies a novel genus-specific CYP51 function. New Phytologist. 198(3), 821–835. doi: 10.1111/nph.12193.

Fernandez, J. and Orth, K. (2018). Rise of a cereal killer: The biology of *Magnaporthe oryzae* biotrophic growth. Trends in Microbiology. 26(7), 582–597. doi: 10.1016/j.tim.2017.12.007.

Gao, Q., Zhao, M., Li, F., Gau, Q., Xing, S. and Wang, W. (2008). Expansins and coleoptile elongation in wheat. Protoplasma. 233: 73–81. doi: 10.1007/s00709-008-0303-1.

Gardiner, D. M., Kazan, K. and Manners, J. M. (2009). Nutrient profiling reveals potent inducers of trichothecene biosynthesis in *Fusarium graminearum*. Fungal Genetics and Biology. 46, 604– 613. doi: 10.1016/j.fgb.2009.04.004.

Gardiner, D. M., Osbourne, S., Kazan, K. and Manners, J. M. (2009). Low pH regulates the productions of deoxynivalenol by *Fusarium graminearum*. Microbiology. 155(9), 3149–3156. doi: 10.1099/mic.0.029546-0.

Guenther, J. C. and Trail, F. (2005). The development and differentiation of *Gibberella zeae* (anamorph: *Fusarium graminearum*) during colonization of wheat. Mycologia. 97(1), 229–237. doi: 10.1080/15572536.2006.11832856.

Hallen-Adams, H.E., Wenner, N., Kuldau, G. A. and Trail, F. (2011). Deoxynivalenol biosynthesis-related gene expression during wheat kernel colonization by *Fusarium graminearum*. Phytopathology. 101(9): 1091–1096. doi: 10.1094/PHYTO-01-11-0023.

Hohn, T. M., McCormick, S. P. and Desjardins, A. E. (1993). Evidence for a gene cluster involving trichothecene-pathway biosynthetic genes in *Fusarium sporotrichioides*. Current Genetics. 24, 291–295. doi: 10.1007/bf00336778.

Ilgen, P., Hadeler, B., Maier, F. J. and Schäfer, W. (2009). Developing Kernel and Rachis Node Induce the Trichothecene Pathway *of Fusarium graminearum* During Wheat Head Infection. Molecular Plant Microbe Interactions, 22(8), 899–908. doi: 10.1094/MPMI-22-8-0899.

Jansen, C., von Wettstein, D., Schäfer, W., Kogel, K-H., Felk, A., and Maier, F. J. (2005). Infection patterns in barley and wheat spikes inoculated with wild-type and trichodiene synthase gene disrupted *Fusarium graminearum*. PNAS. 102(46), 16892–16897. Doi: 10.1073/pnas.0508467102.

Kashyap, A., Planas-Marquès, M., Capellades, M., Valls, M. and Coll, N. S. (2021). Blocking intruders: inducible physico-chemical barriers against plant vascular wilt pathogens. Journal of Experimental Botany. 72(2): 184–198. doi: 10.1093/jxb/eraa444.

Lee, J. and Lu, H. (2011). Plasmodesmata: the battleground against intruders. Trends in Plant Science. 16(4): 201–210. doi: 10.1016/j.tplants.2011.01.004.

Lemmens, M., Scholz, U., Berthiller, F., Dall’Asta, C., Koutnik, A., Schuhmacher, R. et al. (2005). The ability to detoxify the mycotoxin deoxynivalenol colocalizes with a major quantitative trait locus for Fusarium Head Blight resistance in wheat. Molecular Plant Microbe Interactions. 18(12), 1318–1324. doi: 10.1094/MPMI-18-131.

McCormick, S. P., Stanley, A. M., Stover, N. A. and Alexander, N. J. (2011). Trichothecenes: from simple to complex mycotoxins. Toxins. 3(7), 802–814. doi: 10.3390/toxins3070802.

McMullen, M., Bergstrom, G., De Wolf, E., Dill-Macky, R., Hershman, D., Shaner, G., et al. (2012). A unified effort to fight an enemy of wheat and barley: Fusarium head blight. Plant Disease. 96(12), 1712–1728. doi: 10.1094/PDIS-03-12-0291.

McMullen, M., Jones, R. and Gallenburg, D. (1997). Scab of wheat and barley: a re-emerging disease of devastating impact. Plant Disease. 81(12), 1340 – 1348. doi: 10.1094/PDIS.1997.81.12.1340.

Mendiburu, F. and Yaseen, M. (2020). Agricolae: statistical procedures for agricultural research. R package version 1.4.0.

Mesterházy, A. (1995). Types and components of resistance to *Fusarium* head blight of wheat. Plant Breeding. 114, 377–386. doi: 10.1111/j.1439-0523.1995.tb00816.x

Mitra, P.P. and Loque. D. (2014). Histochemical staining of *Arabidopsis thaliana* secondary cell wall elements. Journal of Visualized Experiments. 87: 51381. doi: 10.3791/51381.

Parry, D.W., Jenkinson, P. and McLeod, L. (1995). Fusarium ear blight (scab) in small grain cereals— A review. Plant Pathology. 44, 207–238. doi: doi.org/10.1111/j.1365-3059.1995.tb02773.x.

Pestka, J. J. (2008). Mechanisms of Deoxynivalenol-Induced Gene Expression and Apoptosis. *Food Additives and Contamination Part A: Chemistry, Analysis, Control*, Exposure and Risk Assessment. 25(9): 1128–1140. doi: 10.1080/02652030802056626.

Pestka, J. J. (2010). Deoxynivalenol: mechanisms of action, human exposure, and toxicological relevance. Archives of Toxicology. 84, 663–679. doi: 10.1007/s00204-010-0579-8.

Pritsch, C., Muehlbauer, G. J., Bushnell, W. R., Somers, D. A., and Vance, C. P. (2000). Fungal development and induction of defense response genes during early infection of wheat spikes by *Fusarium graminearum*. Molecular Plant Microbe Interactions. 13(2): 159–169. doi: 10.1094/MPMI.2000.13.2.159.

Proctor, R. H., Hohn, T. M. and McCormick, S. P. (1995). Reduced virulence of *Gibberella zeae* caused by disruption of a trichothecene toxin biosynthetic gene. Molecular Plant Microbe Interactions. 8(4), 593–601. doi: 10.1094/mpmi-8-0593.

Qui, H., Zhao, X., Fang, W., Wu, H., Abubakar, Y. S., Lu, G., et al. (2019). Spatiotemporal nature of *Fusarium graminearum*-wheat coleoptile interactions. Phytopathology Research. 1, article 26. doi: 10.1186/s42483-019-0033-7.

Sager, R. E. and Lee, J. (2018). Plasmodesmata at a glance. Journal of Cell Science. 131(11), jcs209346. doi: 10.1242/jcs.209346.

Sakulkoo, W., Oses-Ruiz M., Garcia E. O., Soanes, D. M., Littlejohn, G. R., Hacker, C. et al. (2018). A single fungal MAP kinase controls plant cell-to-cell invasion by the rice blast fungus. Science. 359(6382), 1399–1403. doi: 10.1126/science.aaq0892.

Shin, S., Torres-Acosta, J.A., Heinen, S.J., McCormick, S., Lemmens, M., Paris, M. P. K., Berthiller, F., Adam, G. and Muehlbauer, G. J. (2012). Transgenic *Arabidopsis thaliana* expressing a barley UDP-glucosyltransferase exhibit resistance to the mycotoxin deoxynivalenol. Journal of Experimental Biology. 63(13): 4731–4740. doi: 10.1093/jxb/ers141.

Tokai, T., Koshino, H., Takahashi-Ando, N., Sato, M., Fujimura, M. and Kimura, M. (2007). *Fusarium Tri4* encodes a key multifunctional cytochrome P450 monooxygenase for four consecutive oxygenation steps in trichothecene biosynthesis. Biochemical and Biophysical Research Communications. 353(2): 412–417. doi: 10.1016/j.bbrc.2006.12.033.

Van Der Plank, J. E. (1963). Plant diseases: epidemics and control. Academic Press, New York.

Vaughan, M., Backhouse, D. and Ponte, E. M. D. (2016). Climate change impacts on the ecology of *Fusarium graminearum* species complex and susceptibility of wheat to Fusarium head blight: a review. World Mycotoxin Journal, 9(5), 685–700. doi: 10.3920/WMJ2016.2053

Wang, Z., Yang, B., Zheng, W., Wang, L., Cai, X., Yang, J. et al. (2022). Recognition of glycoside hydrolase 12 proteins by the immune receptor RXEG1 confers *Fusarium* head blight resistance in wheat. Plant Biotechnology Journal. 21(4): 769–781. doi: 10.1111/pbi.13995.

Wanjiru, W. M., Zhensheng, K., and Buchenauer, H. (2002). Importance of cell wall degrading enzymes produced by *Fusarium graminearum* during infection of wheat heads. European Journal of Plant Pathology. 108: 803–810. doi: 10.1023/A:1020847216155.

Wickham, H. (2016). *ggplot2: Elegant Graphics for Data Analysis*. Springer-Verlag, New York.

Wilson, R. A. and Talbot, N. J. (2009). Under pressure: investigating the biology of plant infection by *Magnaporthe oryzae*. Nature Reviews Microbiology, 7, 185–195. doi: 10.1038/nrmicro2032.

Wu, S., Kumar, R., Iswanto, A. B. B, and Kim, J. (2018). Callose balancing at plasmodesmata. Journal of Experimental Botany. 69(22), 5325–5339. doi: 10.1093/jxb/ery317.

Zhang, X-W., Jia, L-J., Zhang, Y., Li, X., Zhang, D. and Tang, W-H. (2012). In planta stage-specific fungal gene profiling elucidates the molecular strategies of *Fusarium graminearum* growing inside wheat coleoptiles. Plant Cell. 24(12). 5159–5176. doi: 10.1105/tpc.112.105957.

