## Supplementary for "The trichothecene mycotoxin deoxynivalenol facilitates cell-to-cell invasion during wheat-tissue colonisation by *Fusarium graminearum*"

**S1 Determination of the DON concentration to be used in point inoculations with *F. graminearum* conidia**

To determine the suitable concentration of DON for inoculum supplementation, a pilot experiment was conducted to demonstrate a lack of toxicity to *F. graminearum* at various concentrations ranging from 0.5 - 10 times concentrations used *in planta*. The 1.5ml microcentrifuge tubes containing 1ml YPD (Formedium Ltd, UK) were inoculated to a concentration of 5x10^4 spores/ml and incubated at 28°C for 24 hours with rotation at 180 rpm. Hyphal growth was measured at 6, 12, 18 and 24 hours after inoculation by light microscopy and analysed on Fiji image processing software (v. 2.3.0) (Schindelin et al., 2012). The concentration selected for the *in planta* experiment (35ppm DON) was considered to be non-detrimental to spore germination or early spore germling growth.

**S2 Toxicity of DON on plant tissue**

Due to the inability to easily detect deoxynivalenol (DON; chemotype 15-ADON) in plant tissues once glycosylation has occurred,  and the lack of macroscopic necrosis observed in wheat spikelets post-inoculation with water containing 35 ppm DON, we set out to confirm the potent toxicity of the DON chemistry (sourced from Sigma-Aldrich, USA) on plant tissues. We utilised an assay described by Shin et al. (2012) and DON concentrations of 0mg/l, 10mg/l (10ppm, 33µM) and 20mg/l (20ppm, 67µM) to produce comparable phenotypes on *Arabidopsis thaliana* (ecotype Col-0; NASC, UK)  during germination and early growth. Briefly, Col-0 seeds were surface sterilised and germinated on the surface of  ½ MS 1% (w/v) agar plates (as described in Armer et al., 2024), with each plate supplemented with a different DON concentration or no DON. We found significant reductions in both the speed of germination and subsequent seedling growth with increasing concentrations of DON, with a notable absence of root proliferation when DON was present, demonstrating its toxicity to plants.

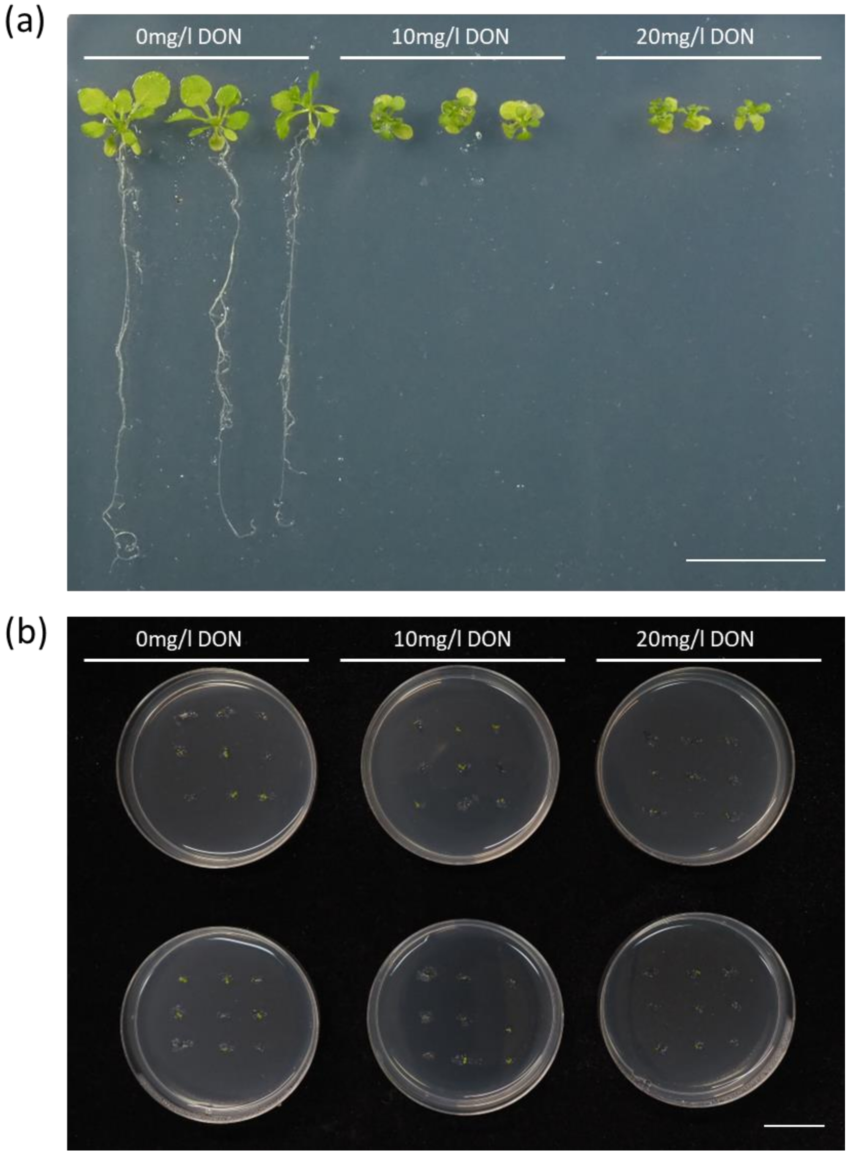


**Demonstration of DON toxicity on plant material.** (a) Growth of *Arabidopsis thaliana* at 3 weeks on ½ MS agar supplemented with 0mg/l DON, 10mg/l DON and 20mg/l DON. (b) Delayed germination of *A. thaliana* on ½ MS agar supplemented with DON. Scale bar = 20mm.

**S3 Primers used in this study (5’-3’)**

| Target | Sense (LP) | Anti-sense (RP) | Use |
|---|---|---|---|

| | | | |
|---|---|---|---|
| FgActin | ATGGTGTCACTCACGTTGTCC | CAGTGGTGGAGAAGGTGTAACC | For RNA expression in wheat coleoptiles, determined by qPCR of cDNA. |
| TRI5 | TCCGTAGCACTATGGACTTTTT | TAGGATGGGCTTCTGAGCCT | |

**S4 Perithecia formation *in vitro* does not require DON**

In natural floral infections in the wheat fields of North America, the typical scab disease symptoms are caused by perithecial development during the later stages of crop maturation (Guenther and Trail, 2004). To determine whether the *ΔTri5* mutant could successfully sexually reproduce, and thereby determine if DON is required for this process, perithecia were induced *in vitro* using a highly reproducible method. The *ΔTri5* mutant was able to produce abundant perithecia that were macroscopically indistinguishable in size to those produced by WT PH-1. Furthermore, to test the viability of formed perithecia, successful ascospore discharge from perithecia was observed by light microscopy for both the WT and *ΔTri5 F. graminearum* strains.

**S5 Cell wall thickness of adaxial cell layer in resin samples.**

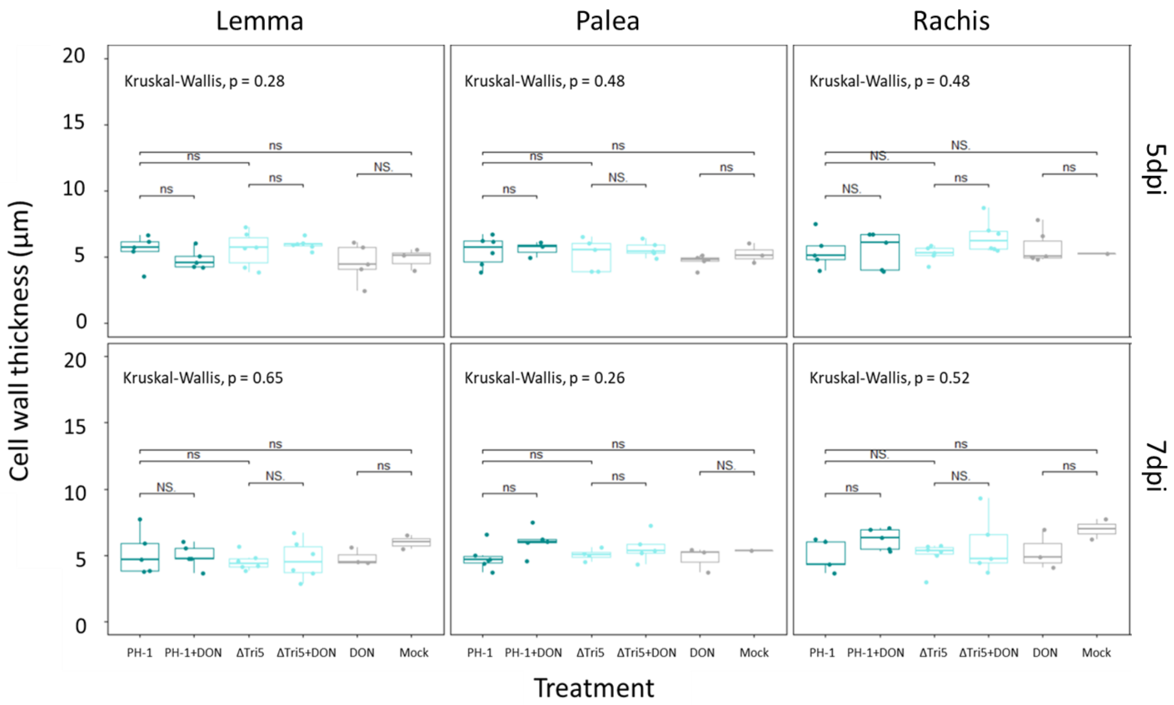


Wheat spikelet tissues of palea, lemma and rachis at 5 and 7dpi time points were analysed, with an average of 10 measurements from a representative resin image of each biological replicate analysed. No significance was determined by Kruskal-Wallis.

**S6 Immunofluorescence detection of callose in sectioned floral tissues**

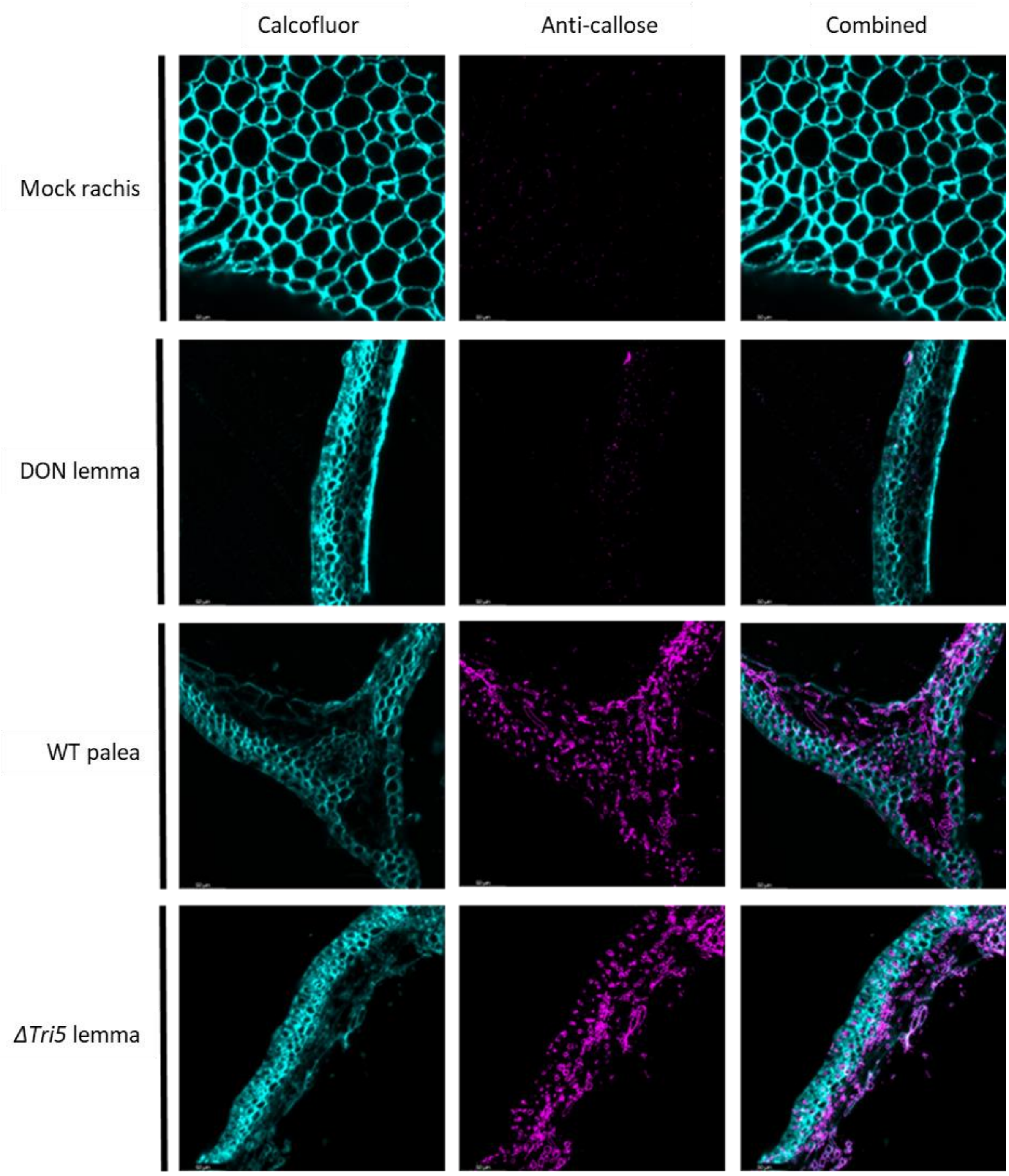


**Immunofluorescence detection of callose in sectioned floral tissues.** Wheat floral tissues from palea, lemma and rachis embedded in LR white resin and sectioned to 1µm then immuno-labelled with anti-callose antibodies and secondarily conjugated with AlexaFluor-488 (magenta). Wheat cell walls are counterstained with calcofluor white (cyan). Selected images of Control, PH-1 (WT) , DON, and *ΔTri5*-infected wheat floral tissues at 5dpi demonstrating typical observations of each interaction.

**S7 Quantification of immuno-labelled callose deposits in wheat spikelet resin sections**

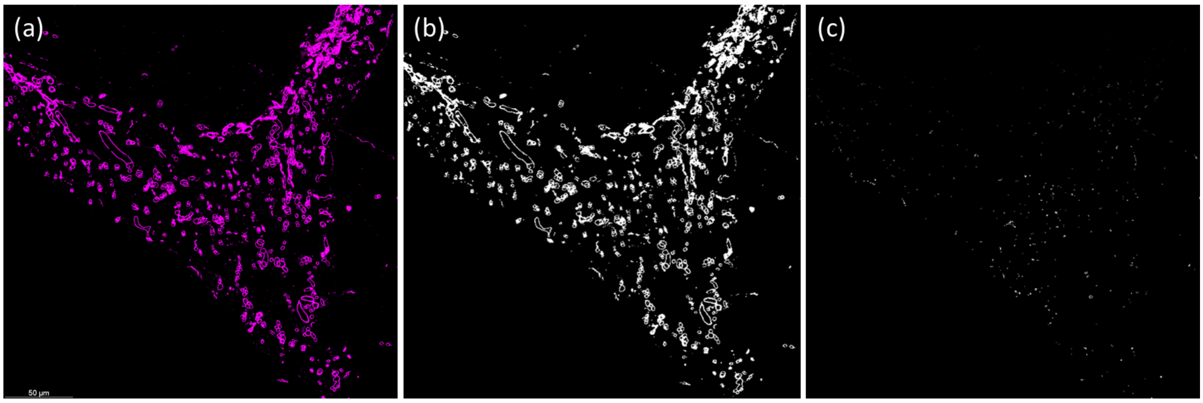


**Example methodology for the quantification of immunofluorescence detection of callose in sectioned floral tissues.** Wheat palea tissue infected with the wildtype PH-1 strain of *F. graminearum* at 5dpi and immuno-labelled for callose, with the secondary antibody conjugated to the fluorophore Alexa Fluor-488. (a) the RGB channel for the emission spectra 510nm- 530nm to detect callose (magenta). β-1,3-glucans in the fungal cell wall have cross-reactivity with the anti-callose antibody. (b) RGB images are converted into binary masks for measurement of particles. (c) Using the Analyse Particles tool in Fiji, particles are counted between 2-19 pixel units in size. This reduces counts due to noise (1 pixel in size) but eliminates pixels attributed to β-1,3-glucans in the fungal cell wall.

**S8 Phloroglucinol staining of infected spikelets for the detection of lignin.**

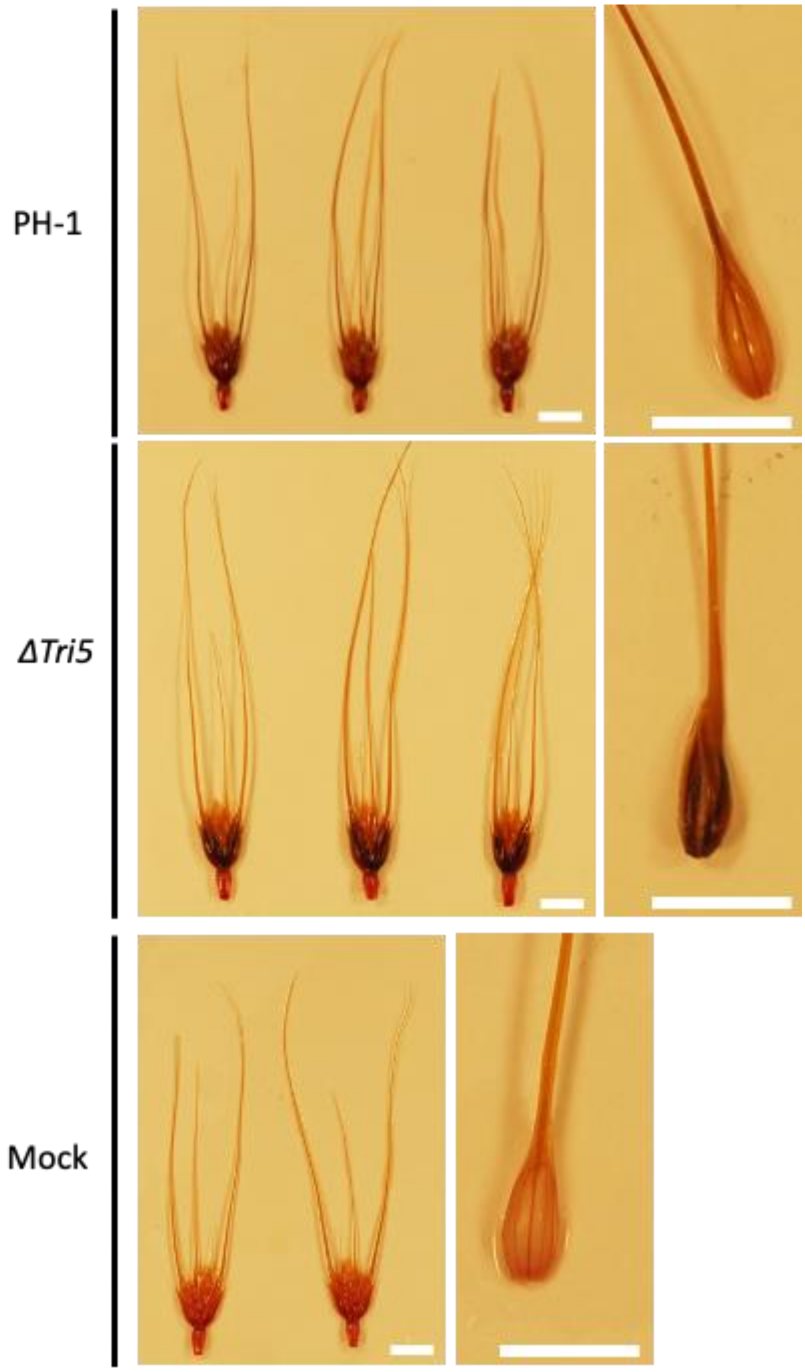


Darker staining of the tissues indicates a greater quantity of lignin. (a) PH-1 - infected spikelet, (b) *ΔTri5*-infected spikelet, (c) Mock-inoculated spikelet. Spikelet component tissues: Lemma demonstrated an increase in phloroglucinol staining component, shown to the left of each treatment, indicating an increase in lignin content. N.B. Point inoculations occur between the lemma and palea tissues. All spikelets were collected at 5dpi and are of the wheat cv. Apogee. Scale bar = 10mm.
